# Beyond the Genotype: A Multi-Omic Analysis of APOEe4’s Role in Alzheimer’s Disease

**DOI:** 10.1101/2025.10.16.682426

**Authors:** Yaroslav Markov, Ahana Priyanka, Leqi Xu, Weiwei Wang, Kyra Thrush-Evensen, John Gonzalez, Daniel Borrus, Jessica Kasamoto, Raghav Sehgal, Grace Zou, Jenel Fraij, Becky C. Carlyle, Steve Horvath, David A Bennett, Hongyu Zhao, Christopher H. van Dyck, TuKiet T Lam, Morgan E. Levine, Albert T. Higgins-Chen

## Abstract

Alzheimer’s disease (AD) is characterized by widespread molecular dysregulation, with the APOEe4 allele recognized as its strongest genetic risk factor. However, the mechanisms by which APOEe4 drives distinct molecular changes – whether by exacerbating pathology or triggering compensatory responses – remain incompletely understood. We generated and analyzed proteomic, epigenetic, and genetic data from post-mortem dorsolateral prefrontal cortex samples of a uniquely APOEe4-enriched subset of the Religious Orders Study and Memory and Aging Project (ROSMAP). Specifically, we generated DIA LC-MS proteomic data (n = 302), analyzed previously generated DNA methylation profiles from our group (n = 310), and used published whole-genome sequencing data (n = 254) to compute polygenic risk scores (PRS). In this cohort, 69% (n = 214) were APOEe4 carriers, and 19.6% (n = 42) of them showed no pathological evidence of AD based on NIA-Reagan criteria, enabling identification of APOEe4-related risk and resilience mechanisms.

In the absence of AD, APOEe4 carriers exhibited lower levels of 27 proteins, suggesting early synaptic (e.g., VAMP1, SYN3, CASKIN1) and metabolic (e.g., GLUD1, PI4KA) vulnerability. By contrast, APOEe4 carriers with AD displayed marked upregulation of inflammatory and proteostatic proteins (e.g., GNAO1, AHNAK, FGG, HEBP1, APEX1, RAB4A, SLC12A5, LRP1, BAG6) and hypermethylation of cg06329447 in ELAVL4. Network analyses highlighted convergent disruptions in synaptic transmission, metabolism, and proteostasis – key pathways altered in APOEe4-associated AD. Mediation analyses identified GRIPAP1 and GSTK1 as top protein mediators (accounting for ∼26–33% of APOEe4’s effect), with VAMP1, CASKIN1, DPP3, SYN3, and FGG each contributing ∼9–15%. ELAVL4 hypermethylation also mediated ∼12% of the APOEe4 effect, linking epigenetic dysregulation to disease risk.

To assess whether the identified proteins reflected broader genetic risk for AD or were specific to APOEe4, we calculated PRS both excluding and including the APOE genomic region. While the non-APOE PRS showed no association with identified molecular markers, the APOE-inclusive PRS was significantly associated with eight AD-related proteins in carriers, indicating they are not explained by polygenic risk outside of APOE. Finally, predictive modeling stratified by APOEe4 status revealed that in non-carriers, PRS most effectively classified AD (AUC = 0.73), whereas in carriers, proteomic and epigenetic markers outperformed PRS (AUC up to 0.74).

Together, these findings demonstrate that APOEe4 confers AD risk through early synaptic and metabolic disruptions and later-stage inflammatory and epigenetic changes, laying the groundwork for genotype-tailored biomarker development and therapeutic strategies.

**VISUAL ABSTRACT:** 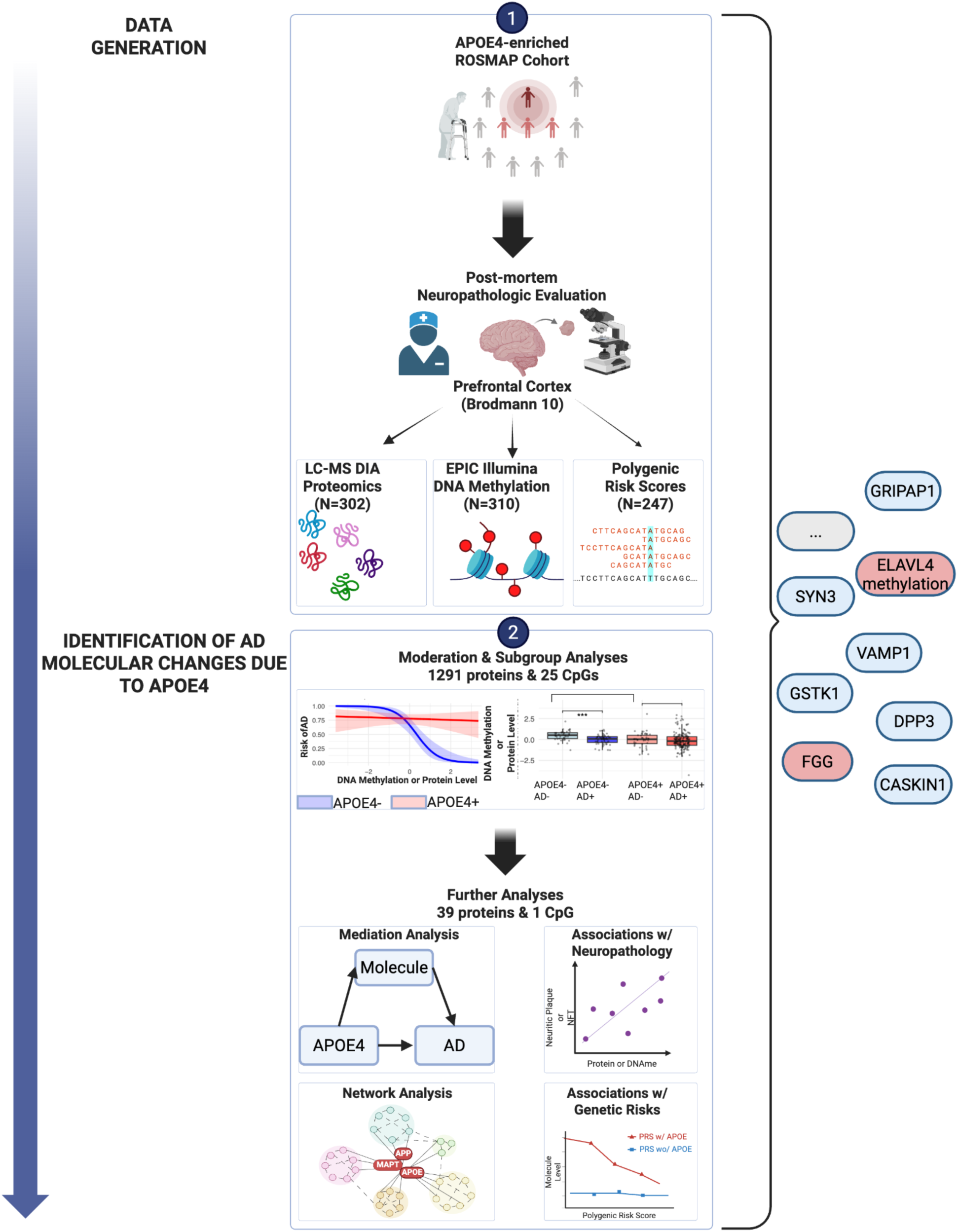

## INTRODUCTION

Alzheimer’s disease (AD) is an escalating global health challenge, currently affecting over 55 million people worldwide, with projections estimating a rise to 139 million by 2050 due to population aging [1]. AD imposes substantial socioeconomic burdens, with global costs exceeding $1.3 trillion annually, driven by long-term care expenditures and productivity losses [2].

Among genetic contributors to late-onset AD, the apolipoprotein E e4 (APOEe4) allele is the most robustly documented risk factor, carried by approximately ∼14% of the global population, with the frequency rising dramatically to ∼40% in patients with AD [3,4]. The allele is associated with a 3-4-fold increased risk (12–15-fold for homozygotes) relative to e3/e3 individuals, as well as with earlier disease onset and more aggressive clinical progression [3,5–8]. Its pathogenic influence extends beyond the classical amyloid–tau mechanisms, disrupting lipid metabolism, mitochondrial function, oxidative stress responses, neuroinflammation, synaptic function, and cerebrovascular integrity [9]; [10–12]. APOEe4 carriers also show earlier reductions of CSF Aβ and elevations of p-tau that precede symptom onset [13–15], suggesting a long preclinical window for molecular intervention. Recent large-scale proteomic work further underscores this point: APOEe4 carriers display a conserved proteomic signature across brain, CSF, and plasma, marked by pro-inflammatory and infection-related immune pathways independent of neurodegenerative diagnosis [16]. Despite these established associations, a detailed molecular understanding of how APOEe4 drives these changes in the brain remains lacking.

Large-scale studies have extensively catalogued protein-level changes associated with AD [17–23]. However, these analyses typically adjust for APOE genotype effects without explicitly investigating genotype-driven biological heterogeneity, obscuring e4-specific changes. Similarly, while epigenetic changes have emerged as critical modulators of gene-environment interactions in neurodegeneration, the impact of APOE genotype on these epigenetic landscapes remains poorly characterized [24–31]. Several CpG sites near genes like ANK1, BIN1, and RHBDF2 consistently associate with AD [32–35], yet few studies examine how APOE genotype might influence these methylation patterns.

To address these knowledge gaps, we leverage large, uniquely structured subset of the Religious Orders Study and Memory and Aging Project (ROSMAP), distinguished by an exceptionally high proportion of APOEe4 carriers (69%, n = 214), with 19.6% (n = 42) of them showing no pathological evidence of AD based on NIA-Reagan criteria. This unique cohort composition provides a rare window into the molecular determinants of both AD risk and resilience in the brain among genetically susceptible populations.

By generating proteomic and DNA methylation data in this cohort, we provide novel multi-omic evidence supporting a biphasic molecular model of APOEe4-driven pathology, characterized by early-stage synaptic and metabolic vulnerabilities that progress to late-stage inflammatory and proteostatic disruptions in clinically manifest AD. Notably, our analyses identify candidate molecular mediators, highlighting potential genotype-specific therapeutic targets to modify risk specifically in APOEe4 carriers.

## RESULTS

### DATASET OVERVIEW

Data for this study were derived from post-mortem prefrontal cortex brain samples obtained from participants in the Religious Orders Study and the Memory and Aging Project (ROSMAP) [36]. All subjects were categorized by both APOE genotype, as described [37], and Alzheimer’s disease (AD) status. AD pathology was assessed post-mortem using a dichotomized version of the National Institute on Aging–Reagan (NIA-Reagan) criteria [38], which combines CERAD plaque scores and Braak neurofibrillary tangle staging, as described [39]. Importantly, these evaluations were performed blinded to clinical data, and classification was based solely on the extent of neuropathology: individuals with intermediate or high levels of pathology were classified as having AD.

The cohort was intentionally enriched for APOEe4 carriers, who made up 69.0% (n=214) of the total sample (**Table 1**). Notably, 19.6% of these carriers (n=42) showed no evidence of AD pathology. Within each APOE genotype group, Non-AD and AD individuals had similar age and sex distributions. This sampling strategy allowed us to specifically examine both risk and resilience mechanisms related to the APOEe4 allele.

**Table 1.**
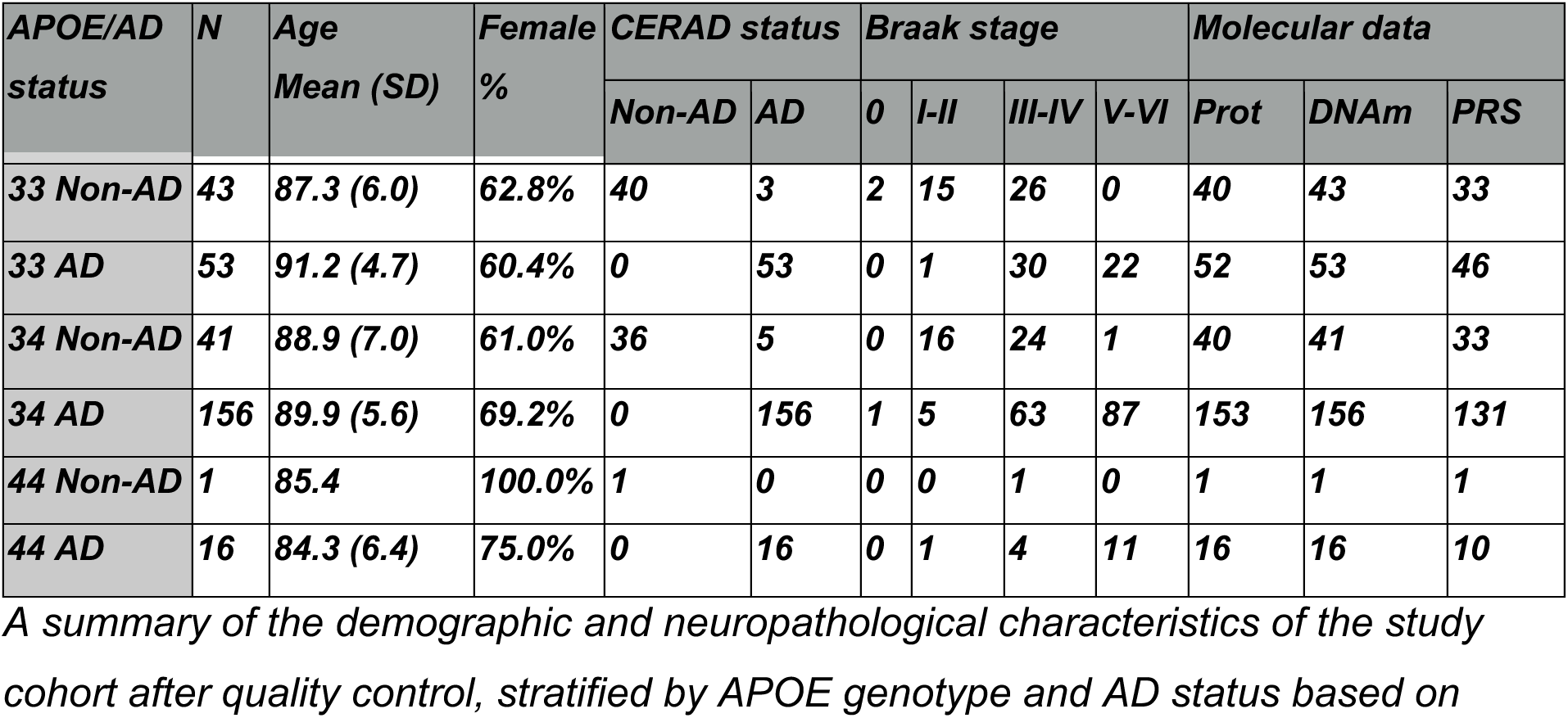
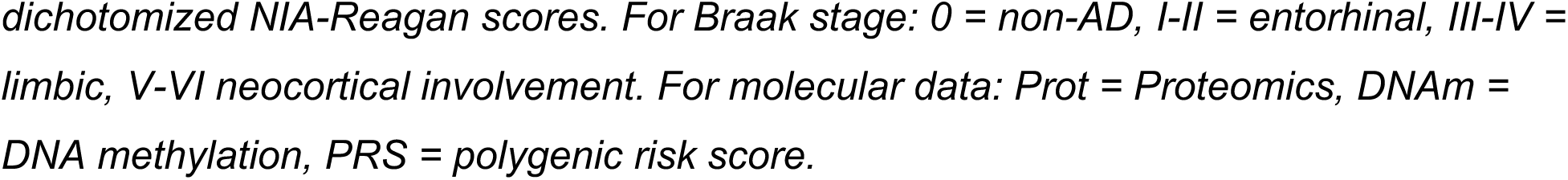

| APOE/AD status | N | Age Mean (SD) | Female % | CERAD status |  | Braak stage |  |  |  | Molecular data |  |  |
| --- | --- | --- | --- | --- | --- | --- | --- | --- | --- | --- | --- | --- |
|  |  |  |  | Non-AD | AD | 0 | I-II | III-IV | V-VI | Prot | DNAm | PRS |
| 33 Non-AD | 43 | 87.3 (6.0) | 62.8% | 40 | 3 | 2 | 15 | 26 | 0 | 40 | 43 | 33 |
| 33 AD | 53 | 91.2 (4.7) | 60.4% | 0 | 53 | 0 | 1 | 30 | 22 | 52 | 53 | 46 |
| 34 Non-AD | 41 | 88.9 (7.0) | 61.0% | 36 | 5 | 0 | 16 | 24 | 1 | 40 | 41 | 33 |
| 34 AD | 156 | 89.9 (5.6) | 69.2% | 0 | 156 | 1 | 5 | 63 | 87 | 153 | 156 | 131 |
| 44 Non-AD | 1 | 85.4 | 100.0% | 1 | 0 | 0 | 0 | 1 | 0 | 1 | 1 | 1 |
| 44 AD | 16 | 84.3 (6.4) | 75.0% | 0 | 16 | 0 | 1 | 4 | 11 | 16 | 16 | 10 |

*A summary of the demographic and neuropathological characteristics of the study cohort after quality control, stratified by APOE genotype and AD status based on*
*dichotomized NIA-Reagan scores. For Braak stage: 0 = non-AD, I-II = entorhinal, III-IV = limbic, V-VI neocortical involvement. For molecular data: Prot = Proteomics, DNAm = DNA methylation, PRS = polygenic risk score.*

Molecular data were collected across all combinations of APOE genotype and AD status, with comparable sampling proportions for proteomics, DNA methylation, and polygenic risk scores (PRS) (**Table 1**). The overlap in data availability across modalities is illustrated in **Supplementary Figure 1**. Proteomic profiling was performed on 302 of the 310 subjects using Liquid Chromatography–Mass Spectrometry (LC-MS) with Data Independent Acquisition (DIA). This approach identified 4,901 unique proteins, including isoforms and post-translationally modified variants, as quantified by Scaffold DIA software. After grouping by UniProt protein names and excluding proteins with >20% missing data, 1,291 proteins were retained for downstream analysis. DNA methylation data were obtained using the Illumina EPIC array on all 310 subjects. PRS were calculated for a subset of 254 individuals using whole-genome sequencing data and effect size weights from the AD GWAS meta-analysis by [40], which included 39,918 AD cases and 358,140 controls.

### BASELINE MOLECULAR CHARACTERISTICS OF ALZHEIMER’S DISEASE

We first sought to characterize baseline molecular alterations associated with Alzheimer’s disease (AD) in our APOEe4-enriched cohort. Differential analyses were performed to compare protein abundance and DNA methylation profiles between individuals with and without AD pathology. All models were adjusted for age, sex, neuronal proportion, and post-mortem interval (PMI) to minimize confounding.

### Proteomics

In the proteomics dataset, differential analysis (FDR ≤ 0.05) revealed 32 proteins significantly upregulated and 16 significantly downregulated in AD (**Figure 1; Supplementary Table 1**). Among the upregulated proteins were canonical AD markers, including amyloid precursor protein (APP; logFC = 2.52, p = 5.34 × 10⁻⁹) and tau (MAPT; logFC = 0.23, p = 1.19 × 10⁻³), alongside less commonly reported proteins such as SLC25A1 (logFC = 0.61, p = 7.71 × 10⁻⁴). Downregulated proteins included key regulators of synaptic transmission (GRIPAP1, logFC = –1.36, p = 6.36 × 10⁻⁶), neuronal stress response (HTT, logFC = –0.74, p = 0.0014), and calcium homeostasis (CSDE1, logFC = –0.98, p = 4.12 × 10⁻⁵) [41]; [42]; [43].

**Figure 1.**
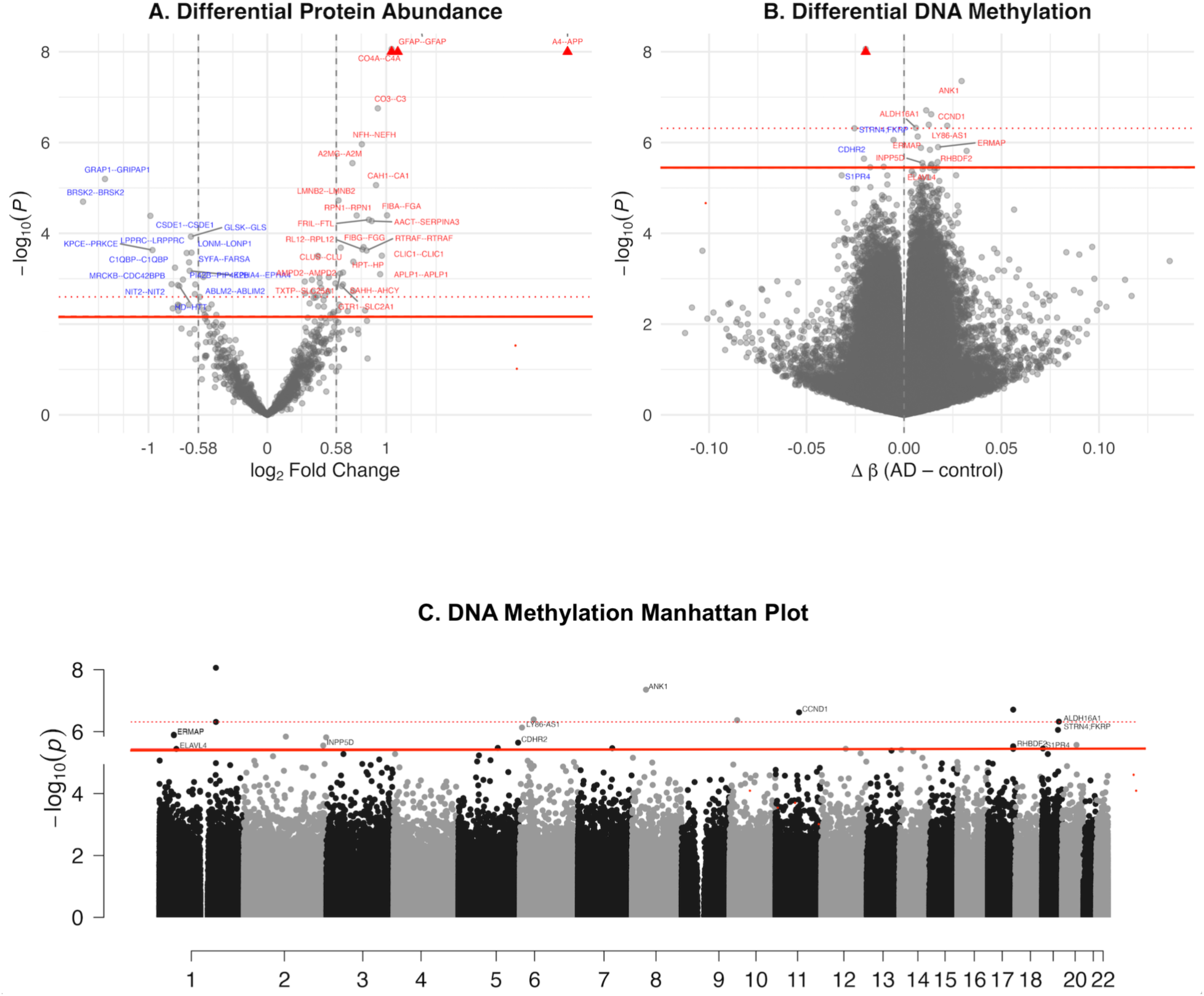
Differential Protein and DNA Methylation Profiles in AD Panel A: Volcano plot of differential protein abundance: log₂ fold change vs. –log₁₀P. Proteins with |log₂FC| ≥ 0.58 and FDR ≤ 0.10 are labeled (red for increased protein level in AD, blue for decreased); points above –log₁₀P = 8 are clipped (red triangles). The dashed horizontal line indicates FDR = 0.10, and the dotted line indicates FDR = 0.05. **Panel B:** Volcano plot of differential DNA methylation: log₂ fold change (Δβ) vs. –log₁₀P. CpGs meeting FDR ≤ 0.10 are labeled (red for increased methylation in AD, blue for decreased). Points above –log₁₀P = 8 are clipped (red triangles). The dashed horizontal line indicates FDR = 0.10, and the dotted line indicates FDR = 0.05. **Panel C:** Manhattan plot of genome-wide differential DNA methylation (AD vs. control) in prefrontal cortex, adjusted for age, sex, neuronal proportion, and post-mortem interval. The dashed horizontal line indicates FDR = 0.10, and the dotted line indicates FDR = 0.05.

To expand the set for enrichment analysis, we relaxed the FDR cutoff to 0.1, yielding 49 upregulated and 27 downregulated proteins. GO enrichment of the upregulated proteins revealed involvement in immune-related processes, filament organization, blood coagulation, and inflammatory response, emphasizing their likely role in AD pathology. For downregulated proteins, the top driver GO terms included synaptic vesicle recycling via endosome, regulation of modification of synaptic structure, positive regulation of macroautophagy, and vesicle-mediated transport in synapse, highlighting disruptions in synaptic maintenance, plasticity, and intracellular trafficking (**Supplementary Figure 2**). Collectively, these results align with established models of AD and reflect coordinated dysregulation across immune, synaptic, and structural pathways.

### DNA Methylation

Differential methylation analysis in prefrontal cortex samples comparing AD and non-AD individuals identified six hypermethylated and two hypomethylated CpG sites at FDR < 0.05 (19 and 6, respectively, at FDR < 0.1) (**Figure 1**; **Supplementary Table 2**). As expected, we observed hypermethylation of ANK1 (cg05066959; ΔBeta = 0.029, p = 4.42 × 10⁻⁸), a well-established epigenetic marker of neurodegeneration [44] implicated in cytoskeletal dysregulation. Several other loci point to novel candidate targets for future epigenetic studies in AD. These included LY86-AS1 (cg06701353; ΔBeta = 0.007, p = 7.37 × 10⁻⁷), an lncRNA inversely associated with Braak stage [45], and two CpG sites in ERMAP (cg25285237, cg12410370; ΔBeta = 0.017, p = 1.26 × 10⁻⁶ and ΔBeta = 0.0086, p = 1.30 × 10⁻⁶), a gene involved in macrophage and T-cell modulation during amyloid clearance [46]. cg06329447 in ELAVL4 (ΔBeta = 0.013, p = 3.60 × 10⁻⁶) was hypermethylated. ELAVL4 is an RNA-binding protein involved in amyloid regulation [47], and its epigenetic alteration may play a role in AD pathogenesis. Hypomethylated sites included CDHR2 (cg00332146; ΔBeta = –0.020, p = 2.26 × 10⁻⁶), which harbors AD-associated exonic variants [48]; S1PR4 (cg17518965; ΔBeta = –0.017, p = 3.50 × 10⁻⁶), potentially reflecting increased immune signaling [49]; and STRN4 (cg04247584; ΔBeta = –0.0054, p = 8.80 × 10⁻⁷), previously implicated in tauopathy mouse models [50].

Taken together, these differentially methylated sites highlight a combination of known and novel epigenetic changes in AD, including immune, cytoskeletal, and RNA-regulatory processes.

### IDENTIFYING MOLECULAR MODERATORS OF THE APOE4 X AD RELATIONSHIP

To better understand how APOE genotype modifies molecular changes in Alzheimer’s disease (AD), we used interaction models to test whether the relationship between AD status and molecular features differs between APOEe4 carriers and non-carriers.

Because our study aimed to uncover molecular factors that may either confer risk or resilience in the context of APOEe4, this analysis was central to our framework. To interpret these interactions, we also categorized individuals into four biologically meaningful genotype–phenotype subgroups: APOEe4−/AD−, APOEe4−/AD+, APOEe4+/AD−, and APOEe4+/AD+ (**Supplementer Table 3**). This allowed us to better characterize molecular signatures that may underlie susceptibility to, or protection from, AD in APOEe4 carriers.

### Proteomics

Our analysis identified 19 proteins with significant APOE × AD interaction effects at FDR < 0.05 (**Supplementary Table 4**). To enable broader biological interpretation and downstream network analyses, we also considered proteins at a more lenient threshold (FDR < 0.1), yielding 121 candidates. We examined how their expression changed across the four genotype–phenotype subgroups (APOEe4−/AD−, APOEe4−/AD+, APOEe4+/AD−, and APOEe4+/AD+) to distinguish between shared disease signatures and genotype-specific effects.

We found that 94 proteins were significantly decreased in AD among APOEe4 non-carriers (**Figure 2**). Within this group, 29 proteins also exhibited lower expression in healthy APOEe4 carriers compared to non-carriers, suggesting that APOEe4 may predispose individuals to a lower baseline of protective proteins that are not further altered by AD itself. This may represent a molecular vulnerability that increases susceptibility to disease. A subset of proteins, including GRIPAP1, BSN, EPHA4, and IDH3A, was consistently downregulated in both carriers and non-carriers with AD, suggesting these changes are robust hallmarks of AD irrespective of genotype. Notably, GRIPAP1 was also reduced in healthy carriers relative to non-carriers. Together, these findings point to both APOEe4-independent and APOEe4-specific mechanisms contributing to AD pathogenesis.

**Figure 2.**
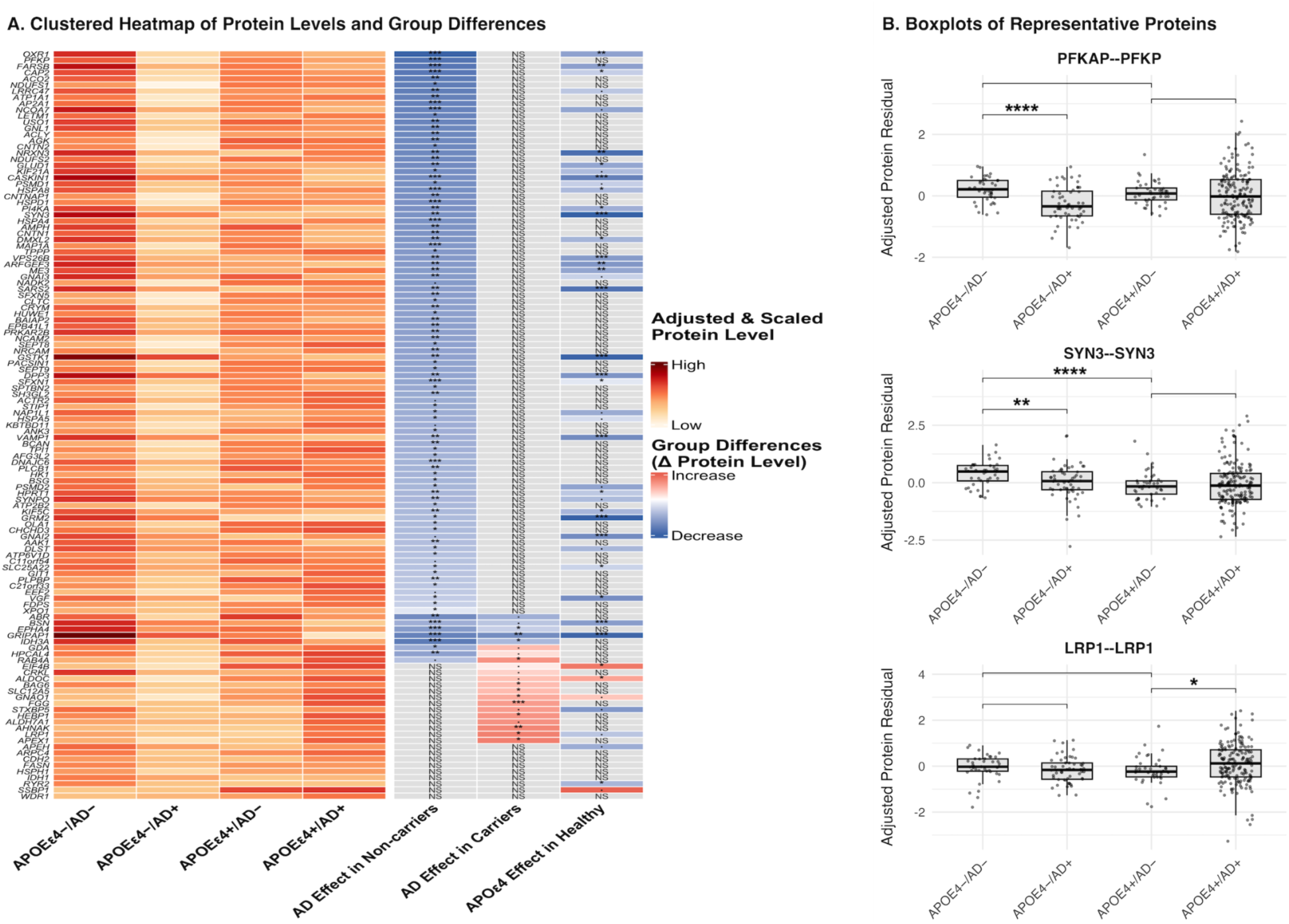
Differential Protein Expression Across APOE and AD Status Groups Panel A: Clustered heatmap displaying scaled, covariate-adjusted mean protein levels for each APOE/AD subgroup: APOE4−/AD−, APOE4−/AD+, APOE4+/AD−, and APOE4+/AD+. In the primary heatmap, colors range from light yellow (lower expression) to red (higher expression). The adjacent contrast heatmap shows directional group-level differences (Δ protein levels) for selected pairwise comparisons. Color indicates direction and magnitude of change: darker blue represents a relative decrease, darker red a relative increase, and grey indicates a nonsignificant comparison. Statistical significance is annotated directly within the contrast heatmap: p< 0.001 (***), p≤ 0.01 (**), p≤ 0.05 (*), p< 0.1 (·), and “NS” for nonsignificant (p ≥ 0.1). **Panel B:** Boxplots of three representative proteins illustrating distinct patterns of differential expression across APOE/AD subgroups. Residual protein levels were adjusted for age, sex, postmortem interval, and neuronal proportion. Each dot represents an individual sample. Statistical comparisons between selected subgroup pairs were performed using Wilcoxon tests. Asterisks indicate significance levels as follows: p < 0.001 (***), p < 0.01 (**), p < 0.05 (*).

We also identified several proteins – GNAO1, AHNAK, FGG, HEBP1, APEX1, RAB4A, SLC12A5, LRP1, CRKL, and BAG6 – that were significantly upregulated only in AD among APOEe4 carriers. These changes may reflect either genotype-specific pathological processes or compensatory responses to disease.

Based on these patterns, we prioritized 39 proteins with APOEe4-specific expression shifts in AD for further investigation. These proteins likely represent mechanisms of risk amplification or resilience specific to APOEe4 carriers.

To determine whether these proteins mediate the effect of APOEe4 on AD risk, we applied mediation analysis. In this context, mediation analysis quantifies how much of APOEe4’s association with AD can be explained by changes in a given protein. Seven proteins showed statistically significant mediation effects (ACME p < 0.05; **Supplementary Table 5**). GRIPAP1 and GSTK1 explained approximately 26–33% of the total APOEe4 effect, suggesting they may play central roles in linking APOEe4 to AD pathology. VAMP1, CASKIN1, DPP3, and SYN3 mediated 9–15% each, with most showing decreased expression in both healthy APOEe4 carriers and AD in non-carriers—consistent with a potential role in early vulnerability. FGG, in contrast, was upregulated in APOEe4 carriers with AD and showed modest mediation (∼10%), indicating it may be part of a different, possibly later-acting disease mechanism. These results highlight potential molecular intermediaries of APOEe4-associated AD risk and offer new targets for further mechanistic and therapeutic investigation.

To link these findings to neuropathological hallmarks, we assessed associations between the 39 prioritized proteins and two standard pathological staging systems: CERAD scores (for amyloid plaques) and Braak stage (for tau tangles). Sixteen proteins were significantly associated with CERAD scores (p < 0.05; **Supplementary Table 6**), with directions consistent with their associations with AD and APOEe4 status. Only six proteins – AHNAK, GNAO1, FGG, KIF5C, GNAI2, and LRP1 – were associated with Braak stage, suggesting that the majority of APOEe4-specific proteins may be more strongly tied to amyloid pathology than tau. This supports the notion that APOEe4 primarily modulates the amyloidogenic pathway and highlights several proteins with potential for early biomarker development or targeted intervention.

### DNA Methylation

Given the high dimensionality of this dataset, we first filtered CpG sites associated with AD at an FDR < 0.1. This preprocessing step reduced the number of sites under consideration and improved power for detecting meaningful APOE × AD interaction effects.

Among the remaining sites, cg06329447 – located in the ELAVL4 gene – stood out due to its increased methylation in AD specifically among APOEe4 carriers (interaction p = 0.03). ELAVL4 is a neuronal RNA-binding protein implicated in amyloid processing and synaptic function [47,51], making it a biologically plausible mediator of APOE-related effects.

To assess whether cg06329447 methylation might lie on the causal pathway between APOEe4 and AD, we applied mediation analysis (as previously done for protein markers). This analysis estimated that methylation at cg06329447 accounted for 11.7% of the total APOEe4 effect on AD risk (ACME = 0.0276, 95% CI: 0.00431–0.0600, p = 0.014), suggesting a modest but significant role in mediating genetic risk.

We then evaluated the relationship between cg06329447 methylation and neuropathological burden. Methylation at this site was strongly associated with CERAD scores (β = 1.59, p = 1.39×10⁻⁵), and to a lesser extent with Braak scores (β = 1.71, p = 0.000206), again indicating a tighter link to amyloid pathology than to tau.

While cg06329447 methylation was not directly correlated with ELAVL4 protein levels, we found a significant APOEe4 × ELAVL4 protein interaction in logistic regression models of AD status (interaction p = 0.015). Specifically, higher ELAVL4 protein levels were protective in APOEe4 non-carriers (log odds = –1.03), but this effect was neutralized in carriers (interaction β = 1.15). ELAVL4 protein also negatively correlated with CERAD scores (β = –0.92, p = 0.0218), but showed no significant association with Braak stage.

### PROTEIN NETWORK OF APOEe4-RELATED AD MOLECULAR INTERACTIONS

To elucidate molecular interactions potentially central to AD pathogenesis, particularly in the context of APOE genotype variation, we constructed a protein interaction network guided by our multi-omic findings. The network was built using known protein–protein interactions from the STRING database [52] and included proteins that exhibited significant APOE-associated changes in our analyses. We also incorporated ELAVL4, given its association with the differentially methylated CpG site cg06329447 and its established role in AD pathology. To contextualize these findings within the broader AD framework, core AD-related proteins such as MAPT, APOE, and APP were included as central reference nodes.

The final network consisted of 43 nodes connected by 36 edges, with an average node degree of 1.67. This reflects tighter clustering than expected by chance (expected number of edges: 15), indicating a significant protein–protein interaction enrichment (p < 1.5×10⁻⁶).

We used Markov Cluster Algorithm [53] to partition the network into functional modules, which we annotated based on known protein functions and Gene Ontology (GO) enrichment terms where applicable (**Figure 3**). The resulting clusters were as follows:

**Figure 3.**
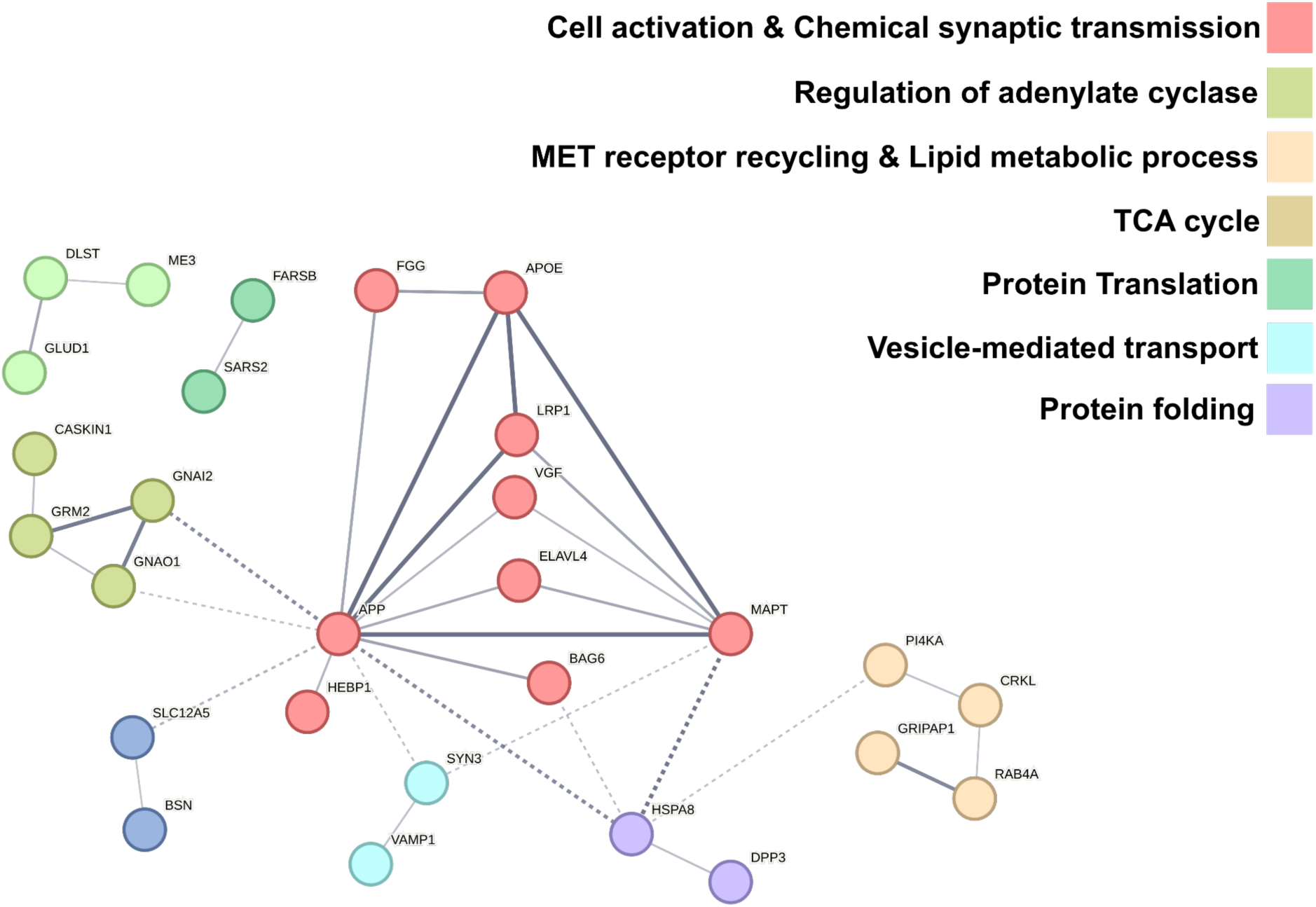
Multi-Omic-Informed Protein Network Anchored by Key AD-Related Genes A network diagram depicting proteins identified through our analyses and literature-based relevance to Alzheimer’s disease. Nodes represent individual proteins and are color-coded according to manually curated functional clusters, including synaptic signaling, metabolic processes, cytoskeletal organization, and immune-related pathways. Key AD-related proteins (MAPT, APOE, and APP) were manually included as reference nodes to anchor the network within the broader context of AD pathogenesis. ELAVL4 was also incorporated due to its relevance to a differentially methylated CpG site (cg06329447) and its established role in amyloid regulation. Solid edges indicate known protein–protein interactions retrieved from the STRING database, with edge thickness reflecting the strength of interaction evidence. Dashed edges represent interactions that connect different functional clusters. Proteins that did not form any connections within the network are not shown.

1) Cell activation + Chemical synaptic transmission:

APP, APOE, BAG6, ELAVL4, FGG, HEBP1, LRP1, MAPT, VGF, SLC12A5, BSN

2) Regulation of adenylate cyclase:

GNAO1, GRM2, GNAI2, CASKIN1

3) MET receptor recycling + Lipid metabolic process:

PI4KA, CRKL, RAB4A, GRIPAP1

4) TCA cycle:

ME3, DLST, GLUD1

5) Protein translation:

FARSB, SARS2

6) Vesicle-mediated transport:

SYN3, VAMP1

7) Protein folding:

HSPA8, DPP3

Fifteen proteins did not connect within the network: NRXN3, OXR1, SLC25A22, HPRT1, AHNAK, ARFGEF3, APEX1, GSTK1, CAP2, LRRC47, SFXN1, VPS26B, DMXL2, SYNPO, KIF5C.

Overall, the network was significantly enriched for multiple GO terms related to synaptic and intracellular communication, including Chemical synaptic transmission, Dendrite development, Synapse assembly and modulation, Behavior, Axo-dendritic transport, Regulation of vesicle-mediated transport – as detailed in **Supplementary Table 7**.

These findings underscore the interconnected nature of synaptic, metabolic, and proteostatic processes modulated by APOE genotype in AD. Notably, HSPA8 – a molecular chaperone involved in protein folding – exhibited high connectivity within the network, linking multiple functional modules through five distinct interactions.

### INTEGRATION OF POLYGENIC RISK SCORES

To further elucidate the genetic underpinnings of the proteomic and epigenetic alterations observed in our study, we calculated Polygenic Risk Scores (PRS) to capture the cumulative impact of common AD-associated variants. We derived two versions of PRS: one excluding the APOE locus (non-APOE PRS) and one encompassing the full genome including the APOE region (APOE PRS). This distinction enabled us to distinguish between genome-wide risk and the APOEe4-driven signal.

In logistic regression models assessing neuropathological burden, a one–standard deviation (SD) increase in the non-APOE PRS was associated with a 79% increase in the odds of high AD pathology by NIA-Reagan criteria (OR = 1.79, 95% CI: 1.53–2.09, p = 1.67×10⁻⁴), a 76% increase for Braak stage (OR = 1.76, 95% CI: 1.42–2.17, p = 0.0073), and a 68% increase for CERAD scores (OR = 1.68, 95% CI: 1.44–1.96, p = 7.78×10⁻⁴). In contrast, the APOE-inclusive PRS exhibited substantially stronger associations: each 1-SD increase was linked to a 117% increase in odds of high NIA-Reagan classification (OR = 2.17, 95% CI: 1.57–3.01, p = 6.01×10⁻⁶), an 89% increase for Braak stage (OR = 1.89, 95% CI: 1.26–2.84, p = 0.0032), and a 116% increase for CERAD score (OR = 2.16, 95% CI: 1.56–3.00, p = 8.45×10⁻⁶). These findings underscore the added explanatory power of APOE-linked variants, particularly in relation to amyloid burden as reflected in CERAD scores.

To assess the relationship between PRS and molecular features, we tested associations between both types of PRS and the differentially expressed proteins and CpG sites identified in earlier analyses. The non-APOE PRS, calculated using both continuous shrinkage (csPRS; [54] and summary statistics-based nonparametric (SDPR; [55] methods, showed no significant associations with either protein or methylation markers after multiple testing correction. This suggests that genome-wide risk outside of APOE may act via pathways not directly captured by proteomic or epigenetic signatures in the prefrontal cortex.

In contrast, the APOE-inclusive PRS, calculated using the same two methods, demonstrated significant associations with the expression of eight proteins previously found to be upregulated in AD among APOEe4 carriers. These included: HEBP1 (csPRS: beta=0.18, FDR=0.0046; SDPR: beta=0.18, FDR=0.0057), LRP1 (csPRS: beta=0.17, FDR=0.0174; SDPR: beta=0.15, FDR=0.0282), FGG (csPRS: beta=0.16, FDR=0.0495; SDPR: beta=0.18, FDR=0.0230), AHNAK (csPRS: beta=0.24, FDR=0.0010; SDPR: beta=0.25, FDR=0.0006), GNAO1 (csPRS: beta=0.18, FDR=0.0021; SDPR: beta=0.16, FDR=0.0057), SLC12A5 (csPRS: beta=0.19, FDR=0.0021; SDPR: beta=0.21, FDR=0.0008), BAG6 (csPRS: beta=0.18, FDR=0.0095; SDPR: beta=0.19, FDR=0.0057), and RAB4A (csPRS: beta=0.13, FDR=0.0513; SDPR: beta=0.16, FDR=0.0088).

These results offer additional evidence that these proteins are tightly linked to APOEe4-specific mechanisms rather than broader polygeneic risk, and validate their relevance in genetically driven AD pathology.

A consolidated summary of the major findings across omic layers is presented in Table 2, which outlines the direction and significance of each molecular association and its relation to APOEe4-specific pathology.

**Table 2.**
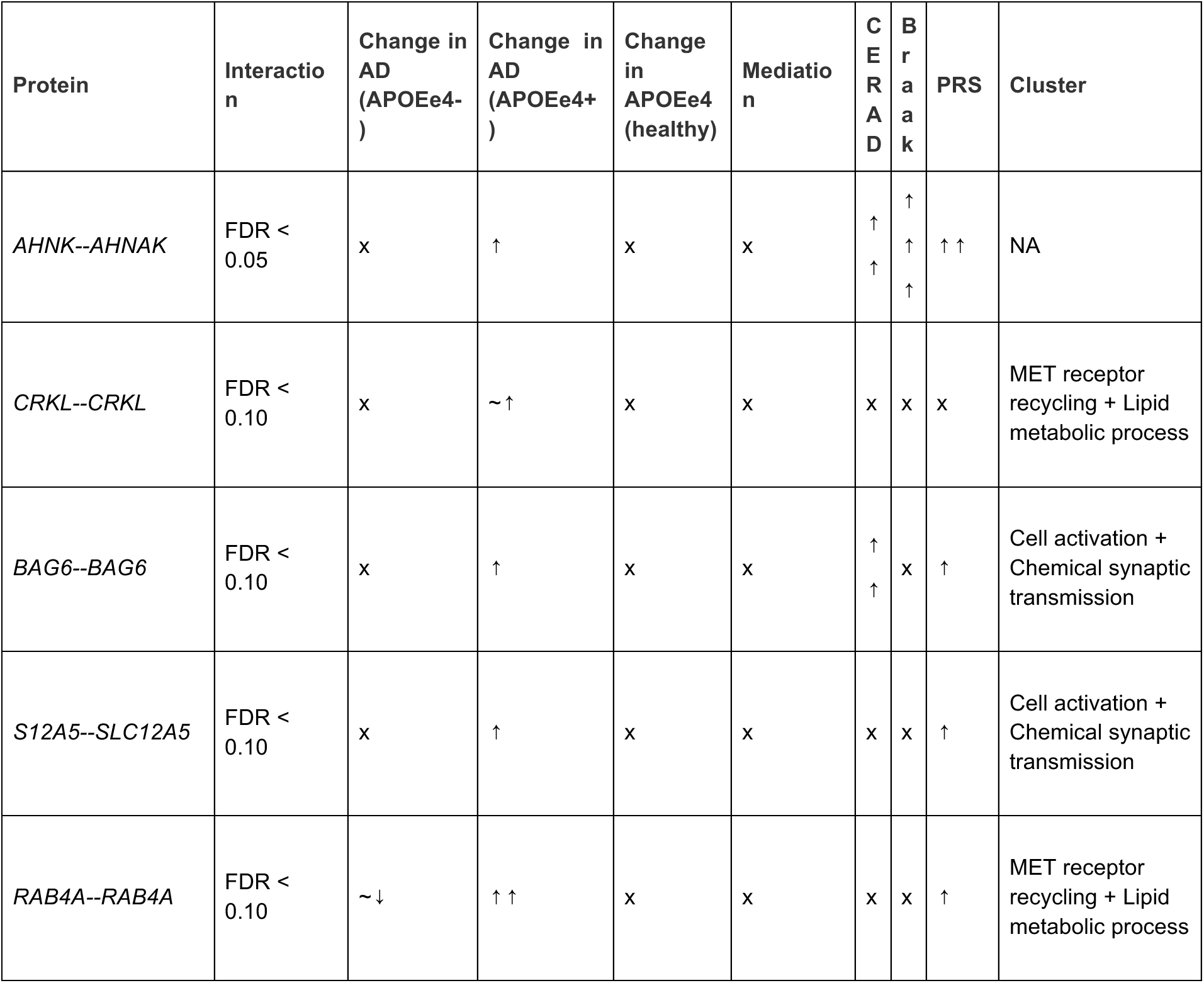
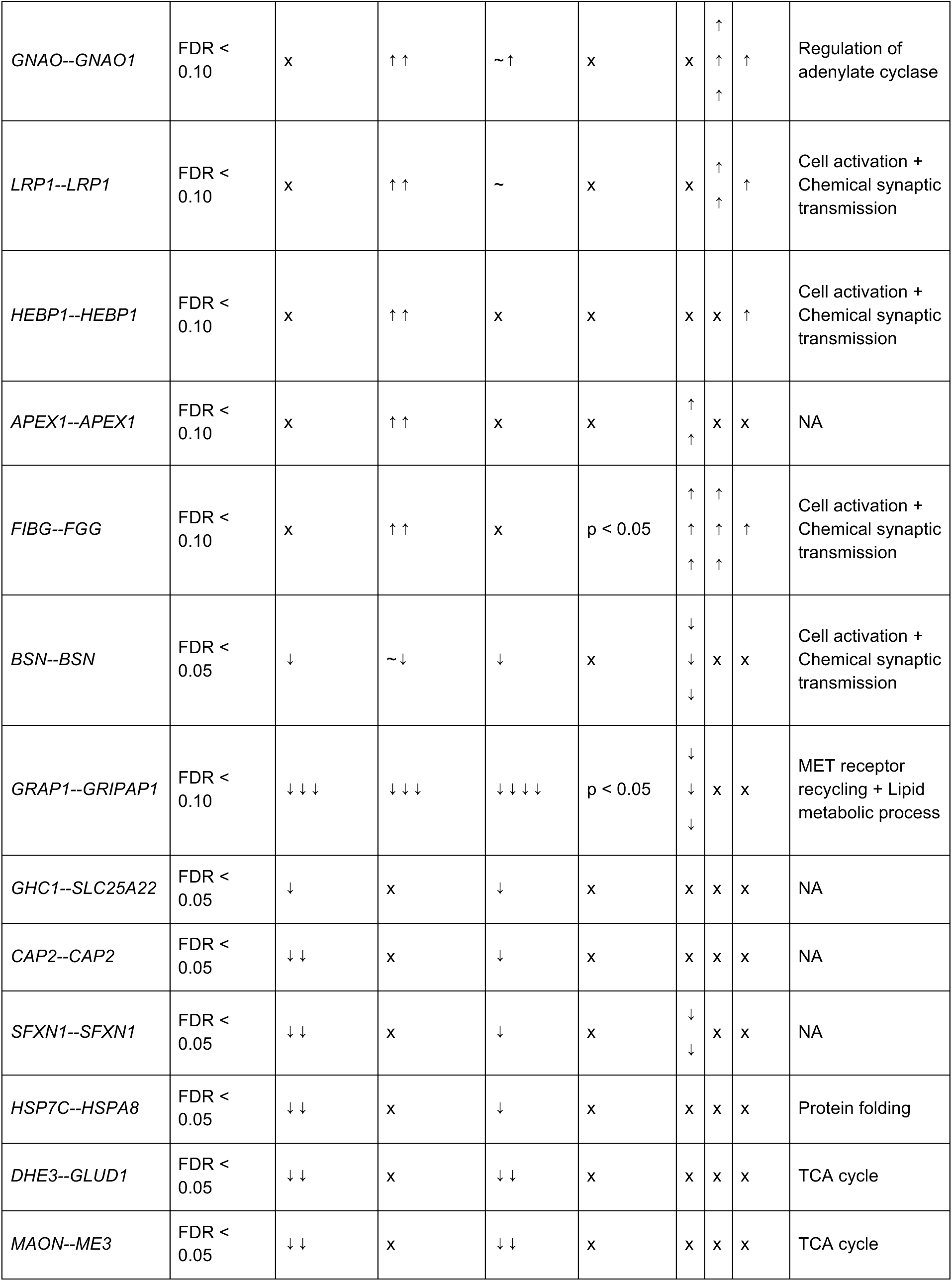
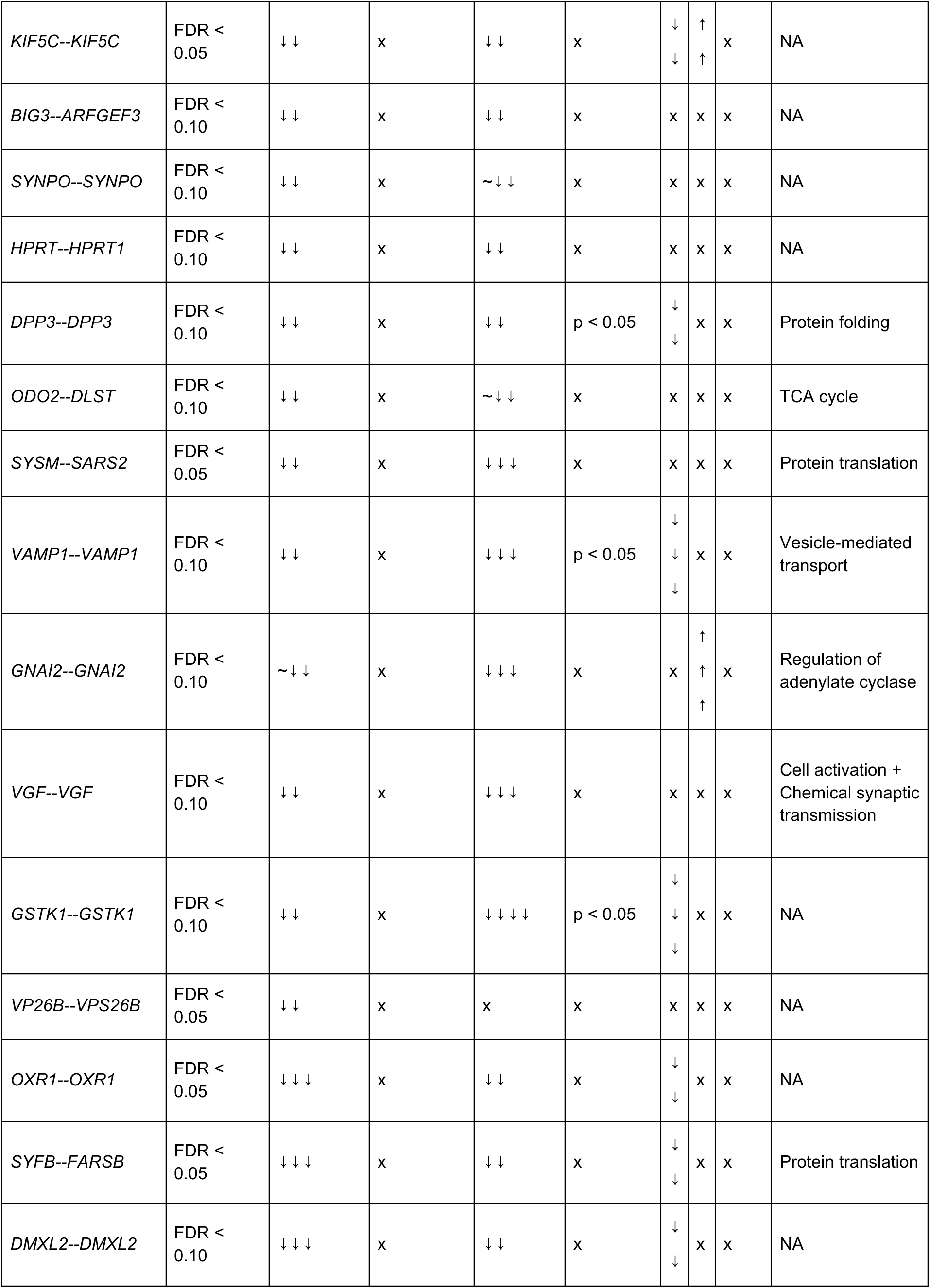
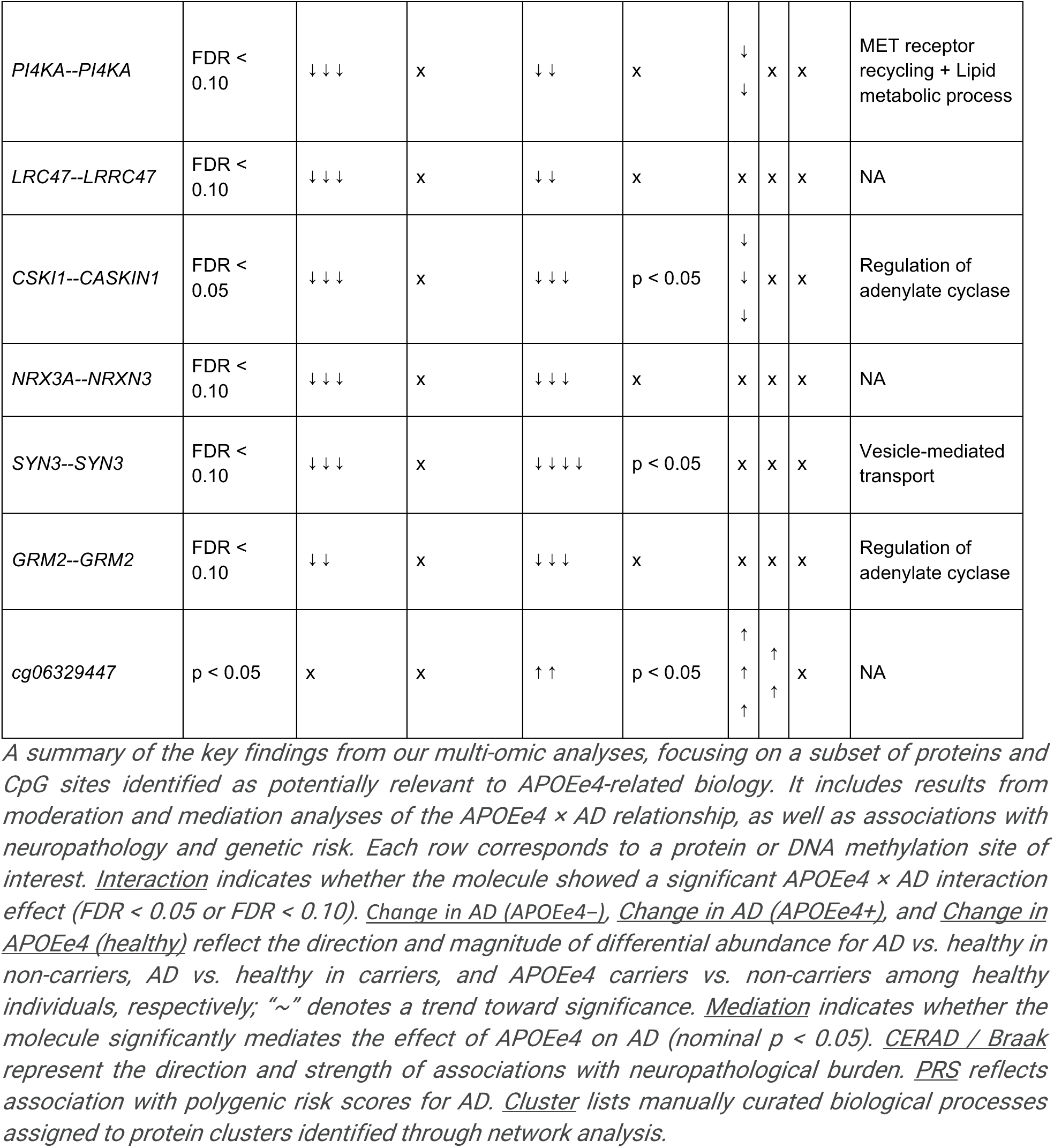

### COMPARISON OF APOEe4-RELATED MOLECULAR CHANGES FOR PREDICTIVE MODELING

Finally, to evaluate the predictive utility of the molecular features identified in our analyses, we assessed how protein expression, DNA methylation, and polygenic risk scores (PRS) performed in classifying AD status. We focused particularly on whether these modalities capture distinct aspects of AD pathology in APOEe4 carriers versus non-carriers.

To summarize proteomic variation, we performed principal component analysis (PCA) separately on each protein cluster defined in our previously constructed interaction network, using only the proteins identified in interaction analyses. The first principal component (PC) from each cluster was used as a representative feature. The 15 proteins that did not cluster were grouped and similarly summarized using PCA. For DNA methylation, we initially examined the predictive power of a single CpG site, cg06329447, which had shown APOEe4-dependent differential methylation.

In APOEe4 non-carriers, predictive modeling yielded an area under the curve (AUC) of 0.69 for proteomic PCs, 0.73 for the APOE-inclusive PRS, and 0.62 for cg06329447 methylation. In contrast, in APOEe4 carriers, the AUC values were 0.63 (proteomics), 0.56 (PRS), and 0.66 (cg06329447 methylation), suggesting that non-genetic markers – particularly DNA methylation – have greater predictive value than PRS in carriers.

Recognizing that a single CpG site may not fully capture the complexity of APOEe4-related epigenetic dysregulation, we next applied weighted gene co-expression network analysis (WGCNA) [56] to 67,654 CpG sites nominally associated with AD (p < 0.05).

This yielded several co-methylation modules whose eigengenes were tested for their predictive utility. Notably, cg06329447 (in ELAVL4) belonged to the pink module, which was strongly associated with AD (β = 0.26, p = 2×10⁻⁵), CERAD score (β = 0.24, p = 2×10⁻⁴), and Braak stage (β = 0.19, p = 0.03). It was also one of two modules that significantly moderated the APOEe4 × AD interaction (interaction coefficient = 0.65, p = 0.03). All ELAVL4-associated CpGs clustered into either the pink or the midnight blue modules.

We then evaluated three sets of predictive models: (1) using the eigengene of the pink module only, (2) using the eigengenes of both the pink and midnight blue modules (representing modules positively associated with AD and that enhance the APOEe4 effect), and (3) using the eigengenes of all four moderating modules, which include the pink and midnight blue modules along with two additional modules – the salmon and light green modules – that were negatively associated with AD and moderated the APOEe4 × AD interaction in the opposite direction (salmon: AD β=–0.42, p=2×10⁻⁴; CERAD β=–0.43, p=2×10⁻⁴; Braak β=–0.30, p=0.05; interaction coefficient=–0.67, p=0.02; light green: AD β=–0.26, p=0.004; CERAD β=–0.25, p=0.007; interaction coefficient=–0.75, p=0.03). Using the eigengene of the pink module alone, the AUC improved to 0.67 in non-carriers and 0.65 in carriers (**Figure 4**). Adding the eigengene of the midnight blue module did not significantly increase performance (AUC = 0.67 in non-carriers, 0.68 in carriers). However, incorporating all four moderating modules – pink, midnight blue, salmon, and light green – raised the AUC to 0.67 for non-carriers and 0.74 for carriers. This suggests that including DNA methylation modules negatively moderating the APOEe4 × AD relationship enhances predictive performance in carriers. Overall, network-based module integration strengthened the predictive value of DNA methylation for both groups.

**Figure 4.**
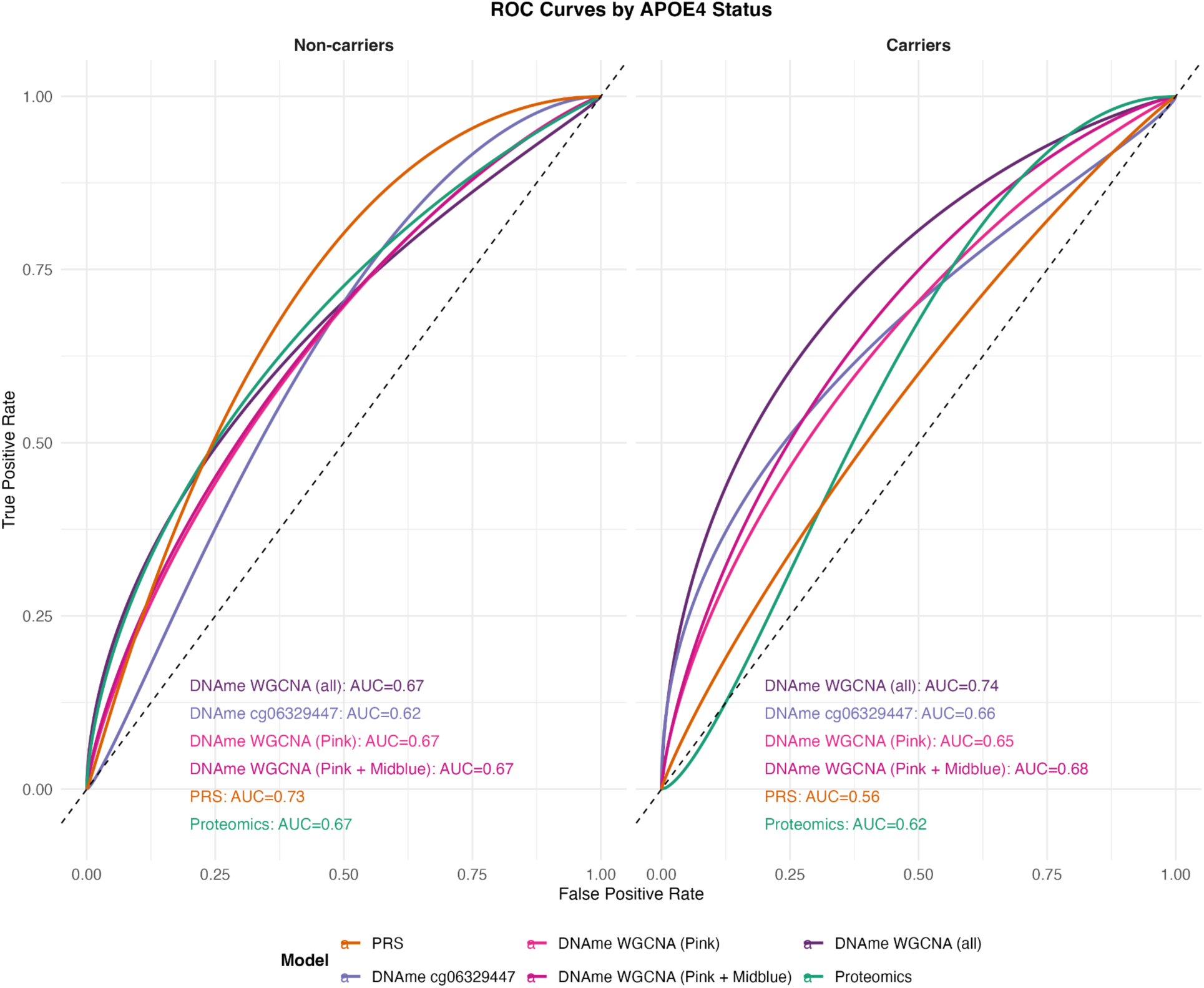
Discriminatory Accuracy of Protein-, PRS-, and Methylation-Based Models in APOEe4 Carriers vs Non-Carriers Receiver operating characteristic (ROC) curves for predictive models stratified by APOEe4 carrier status, showing the area under the curve (AUC) for models built using protein markers, polygenic risk scores (PRS), a single CpG site identified in the interaction analysis, and DNA methylation subnetworks derived from WGCNA. Protein predictors were summarized using principal component analysis (PCA) based on their assigned functional clusters from the network analysis. For methylation-based models, we evaluated three sets of predictors: (1) the eigengene of the pink module alone, (2) eigengenes of the pink and midnight blue modules (positively associated with AD and APOEe4 effect), and (3) eigengenes of all four moderating modules, including the salmon and light green modules, which were negatively associated with AD and moderated the APOEe4 × AD interaction in the opposite direction.

Collectively, our integrative analyses – including interaction modeling, network construction, mediation testing, and predictive modeling – converge to highlight a distinct molecular signature of APOEe4-associated AD. While many changes are shared across genotypes, specific proteins and CpG methylation patterns show genotype-dependent modulation. This work establishes a comprehensive multi-omic framework for understanding APOEe4-related risk and resilience, and identifies molecular candidates for genotype-tailored diagnostic and therapeutic strategies.

## DISCUSSION

In this study, we leveraged a multi-omic framework encompassing proteomic, epigenetic, and genetic data to characterize the molecular landscape associated with the APOEe4 allele in Alzheimer’s disease (AD). By integrating LC-MS proteomics, DNA methylation profiling, and polygenic risk scores (PRS), we identified a two-phase pattern of APOEe4-associated molecular alterations. In cognitively intact APOEe4 carriers, we observed early reductions in proteins related to synaptic function and metabolism, indicating presymptomatic molecular vulnerability. These findings align with recent positron emission tomography imaging studies showing reduced hippocampal and medial temporal lobe synaptic density in APOEe4 carriers prior to clinical symptoms [57]; [58], as well as elevated cerebrospinal fluid levels of the synaptic marker neurogranin in early cognitive impairment [59], underscoring the clinical relevance of our observations. The identified reductions in metabolic enzyme levels are also consistent with previous reports of decreased cerebral glucose metabolism in cognitively healthy APOEe4 carriers [60–64]. In contrast, APOEe4 carriers with AD exhibited increased inflammatory and proteostatic proteins and distinct epigenetic modifications, suggesting a secondary phase involving APOEe4-specific responses to established pathology. This dual-phase model provides a valuable framework for understanding how APOEe4 contributes to early susceptibility and subsequent progression of AD.

### INTERPLAY BETWEEN APOE GENOTYPE AND MOLECULAR CHANGES

Our findings highlight distinct molecular signatures linked to APOEe4 genotype, emphasizing both early vulnerabilities and disease-specific alterations. Even in the absence of clinical AD, APOEe4 carriers displayed reductions in proteins crucial for synaptic integrity and metabolic homeostasis. Synaptic regulators such as VAMP1, essential for synaptic vesicle fusion and Aβ exocytosis [65]; [66], and SYN3, important for dopaminergic neuronal function [67], were notably reduced, aligning with preclinical models demonstrating synaptic deficits associated with APOEe4 [58]. Concurrently, decreased levels of metabolic enzymes GLUD1 and PI4KA, key regulators of glutamate-GABA balance and lipid metabolism [68]; [69], support prior evidence of metabolic dysfunction in APOEe4 carriers [70]. Early oxidative imbalances were suggested by reduced antioxidant proteins OXR1 and DPP3 [71–73]. Mediation analyses identified proteins such as GRIPAP1, GSTK1, VAMP1, SYN3, CASKIN1, and DPP3 as significant intermediaries between APOE genotype and AD, collectively accounting for 10–33% of APOEe4’s total effect, reinforcing the importance of synaptic dysfunction and oxidative stress as mechanisms driving early APOEe4-related vulnerability.

In the context of clinical AD, APOEe4 carriers demonstrated pronounced upregulation of inflammatory, proteostatic, and stress-response pathways, potentially indicative of compensatory responses. Elevated proteins such as HEBP1 and FGG suggest increased neurovascular and coagulation dysfunction [74]; [75], consistent with findings from isogenic iPSC-derived endothelial models showing ApoE4-linked alterations in blood coagulation and barrier integrity pathways [76]. Stress-response proteins including APEX1 and BAG6 further indicated attempts to mitigate proteostatic and cytoskeletal stress [77]. The increased expression of LRP1, a receptor modulated by APOEe4 and involved in Aβ clearance, may represent an inadequate compensatory mechanism [78]. Complementing these proteomic changes, hypermethylation at cg06329447 in ELAVL4, a critical RNA-binding regulator of Aβ production [47,51], emerged as a significant epigenetic alteration in APOEe4 carriers with AD. This hypermethylation mediated approximately 12% of the APOEe4 effect on AD, possibly destabilizing ELAVL4 transcripts and accelerating disease progression. Overall, our data suggest APOEe4 shifts the molecular balance from early synaptic-metabolic deficiencies to late-stage inflammatory and proteostatic stress.

A systems-level interaction network analysis, informed by the STRING database, contextualized these molecular changes into distinct functional clusters, highlighting synaptic transmission, adenylate cyclase signaling, metabolic regulation, vesicle-mediated transport, and proteostasis. A prominent cluster linked synaptic transmission and neuroinflammation (APP, APOE, BAG6, ELAVL4, FGG, HEBP1, LRP1, MAPT, VGF, SLC12A5, BSN), underscoring APOEe4’s role in neurovascular dysfunction and synaptic vulnerability [79]; [58]. Another cluster centered on disrupted adenylate cyclase signaling (GNAO1, GRM2, GNAI2, CASKIN1), critical for synaptic plasticity and cognitive function [80]; [81]; [82]). Metabolic dysfunction involving mitochondrial enzymes GLUD1, DLST, and ME3 further pointed to bioenergetic deficits linked to APOEe4 [68]; [83]. Cellular trafficking disruptions were captured by a module involving MET receptor recycling and lipid metabolism (PI4KA, CRKL, RAB4A, GRIPAP1), aligning with APOEe4-driven lipid dysregulation and impaired receptor trafficking [84];

[85]; [86]; [70]). Vesicle-mediated transport deficits involving SYN3 and VAMP1 highlighted impaired intracellular trafficking related directly to Alzheimer’s pathology. Finally, proteostasis disruption emerged through proteins involved in protein folding and mitigating oxidative stress (HSPA8, DPP3), emphasizing the intersection between redox imbalance and protein quality control mechanisms ([87]; [72]). DPP3 activates neuroprotective NRF2 pathways by competing with KEAP1 binding during oxidative stress [88]; [89]), while HSPA8 is a molecular chaperone decreased in AD [87].

### INTEGRATION OF POLYGENIC RISK SCORES AND PREDICTIVE MODELING

By deriving two separate Polygenic Risk Scores (PRS) – one excluding the APOE region (non-APOE PRS) and one including it (APOE PRS) – we evaluated how genetic risk factors relate to the molecular signatures identified. The non-APOE PRS showed no significant correlations with our proteomic data, indicating that non-APOE genetic risk might act through pathways not captured by prefrontal cortex proteomics. In contrast, the APOE PRS confirmed significant associations with eight proteins increased in AD among APOEe4 carriers: HEBP1, LRP1, FGG, AHNAK, GNAO1, SLC12A5, BAG6, and RAB4A, further supporting their involvement in APOEe4-driven AD pathology.

Predictive modeling using proteomic, epigenetic, and genetic data highlighted distinct molecular signatures in APOEe4 carriers versus non-carriers. In non-carriers, the APOE-inclusive PRS (AUC = 0.73) outperformed non-genetic molecular features, suggesting broader genetic contributions to sporadic AD. In contrast, in APOEe4 carriers, non-genetic markers – particularly WGCNA-integrated epigenetic signatures – demonstrated superior discriminative power compared to the PRS (AUC = 0.56), achieving an AUC of 0.74 when combining multiple methylation modules. Notably, using the eigengene of a single WGCNA module (pink) already improved prediction over individual CpG sites in both groups, with further gains observed in carriers when including additional modules, especially those negatively moderating the APOEe4 × AD interaction (salmon and light green). This suggests that coordinated, network-level methylation changes may better reflect biologically meaningful dysregulation than single-site signals, and that incorporating both risk-and resilience-linked modules enhances model performance in genetically susceptible individuals. Proteomic markers also showed better predictive power in carriers, with an AUC of 0.63. Thus, these findings reinforce the value of omic biomarkers in identifying at-risk individuals and tailoring therapeutic strategies for genetically defined subgroups.

## CONCLUSION/LIMITATIONS

Collectively, our integrated multi-omic analysis delineates a detailed molecular landscape of APOEe4-associated AD pathogenesis, highlighting both the pathogenic and potentially compensatory networks activated by this allele. By merging proteomic, epigenetic, and genetic data, we identified a trajectory in which early synaptic and metabolic deficits predispose APOEe4 carriers to inflammatory, proteomic, and epigenetic dysregulation. These insights lay a promising foundation for the development of genotype-specific biomarkers and personalized therapeutic interventions.

Nonetheless, several limitations must be acknowledged. The cross-sectional, post-mortem design of our study limits causal inference, emphasizing the need for longitudinal investigations – ideally incorporating peripheral biomarkers – to capture dynamic molecular changes in living individuals. In addition, the use of bulk tissue may mask cell-type–specific effects, suggesting that single-cell or spatial approaches could further refine our understanding of neuronal versus glial contributions to APOEe4-associated risk and resilience. Functional validation in APOEe4-relevant cellular or animal models remains essential to firmly establish causal mechanisms, while expanding cohort sizes and including participants from diverse genetic backgrounds will enhance the generalizability of our findings. Despite these challenges, our work advances the field by setting the stage for future studies that not only address current limitations but also pave the way toward more effective, personalized strategies for early diagnosis and intervention.

## METHODS

### Study Population and Tissue Acquisition

Frozen brain tissue samples were obtained from participants in the Religious Orders Study and the Rush Memory and Aging Project (ROSMAP) [36]. All participants enrolled without known dementia and agreed to detailed clinical evaluation and brain donation at death. All studies were approved by an Institutional Review Board of Rush University Medical Center. Each participant signed informed and repository consents and all ROSMAP participants signed an Anatomic Gift Act. For this study, tissue was isolated from Brodmann Area 10 (prefrontal cortex). Sample phenotype data, including clinical diagnoses and neuropathologic assessments, were provided by the Rush University Alzheimer’s Disease Center in accordance with previous publications [90,91].

Alzheimer’s disease (AD) neuropathology was classified based on dichotomized NIA-Reagan diagnostic criteria, which integrate assessments of neurofibrillary tangles and neuritic plaques. Subjects were categorized as having AD pathology if they met the criteria for high or intermediate likelihood and as not having AD if they had low likelihood or no AD pathology. Braak stage was dichotomized for logistic regression analyses into two groups: stages 0-II versus III and above, where stages I-II indicate neurofibrillary tangles predominantly confined to the entorhinal region, stages III-IV indicate limbic region involvement (including the hippocampus), and stages V-VI indicate moderate to severe neocortical involvement. Similarly, CERAD scores were dichotomized, with a diagnosis of AD requiring moderate (probable AD) or frequent neuritic plaques (definite AD) in one or more neocortical regions [39,91].

### Proteomic Data Acquisition and Processing

#### Sample Preparation for LC–MS/MS

Frozen tissues were lysed with a probe sonicator in solubilization buffer (RIPA buffer with serine/threonine protease and phosphatase inhibitors). Lysate was centrifuged at 14.5 x 103 g for 10 minutes at 4 °C in a tabletop centrifuge to pellet cellular debris, and supernatant was collected and protein precipitated with a standard methanol:chloroform:water method. Protein pellets were washed three times with cold MeOH prior to being resuspended in 80uL solubilization buffer (8M urea in 0.4M ammonium bicarbonate). Cysteine of proteins were reduced with 8 µL of 45 mM dithiothreitol (DTT) and incubated at 37 °C for 30 min, then were alkylated with 8 µL of 100 mM iodoacetamide (IAN) and incubated in the dark at room temperature for 30 min. Protein solution was then diluted with water to bring urea concentration to 2 M. Proteins were then digested with sequencing-grade trypsin (Promega, Madison, WI, USA) at a weight ratio of 1:50 (trypsin/protein). Digestion was allowed to occur at 37 °C overnight and more trypsin was added and incubated at 37°C for additional 4 hours. The digestion was quenched with by adding 0.1% formic acid; and peptide mixture was desalted using C18 spin columns (The Nest Group, Inc., Southborough, MA, USA). Eluted peptides were dried in a rotary evaporator and stored until ready for LC MS/MS data collection. The samples were resuspended in 0.2% trifluoroacetic acid (TFA) and 2% acetonitrile (ACN) in water prior to LC–MS/MS analysis.

#### Mass Spectral Data Collection: Data-Independent Acquisition (DIA)

DIA LC–MS/MS was performed using a Waters ACQUITY UPLC M-Class system (Waters Corporation, Milford, MA, USA) coupled to a Q-Exactive HFX (ThermoFisher Scientific, San Jose, CA, USA) mass spectrometer. After injection, the samples were loaded onto a trapping column (nanoEase M/Z Symmetry RP C18 Trap column, 180 µm × 20 mm) at a flow rate of 10 µL/min and separated with a RP C18 column (nanoEase M/Z column Peptide BEH C18, 75 µm × 250 mm). The compositions of mobile phases A and B were 0.1% formic acid in water and 0.1% formic acid in ACN, respectively. The peptides were separated and eluted with a gradient extending from 6% to 25% mobile phase B in 98 min and then to 85% mobile phase B in additional 5 min at a flow rate of 300 nL/min and a column temperature of 37 °C. Column regeneration and up to three blank injections were carried out in between each sample injection to ensure no-carry over. The data were acquired with the mass spectrometer operating in a DIA mode with an isolation window width of 10 Th. The full scan was performed in the range of 400– 1,000 m/z with “Use Quadrupole Isolation” enabled at an Orbitrap resolution of 30,000 at 200 m/z and automatic gain control (AGC) target value of 3 × 106. Fragment ions from each DIA window (e.g., MS2 fragmentation) were generated in the C-trap with higher-energy collision dissociation (HCD) at a normalized collision energy of 28% and detected in the Orbitrap at a resolution of 15,000.

DIA spectra were searched against a Homo Sapiens brain proteome fractionated spectral library generated from DDA LC MS/MS spectra (collected from the same Q-Exactive HFX mass spectrometer) using Scaffold DIA software v. 2.1.0 (Proteome Software, Portland, OR, USA). Within Scaffold DIA, raw files were first converted to the mzML format using ProteoWizard v. 3.0.11748. The samples were then aligned by retention time and individually searched with a mass tolerance of 10 ppm for the precursor ions and a fragment mass tolerance of 10 ppm. The data acquisition type was set to “Overlapping margins of 2 Da”, and the maximum missed cleavages was set to 2. Peptides with charge states between 2 and 4 and 6-30 amino acids in length were considered for quantitation, and the resulting peptides were filtered by Percolator v. 3.01 at a threshold FDR of 0.01. Peptide quantification was performed by EncyclopeDIA v. 0.9.2 and five of the highest quality fragment ions were selected for protein quantitation. Proteins containing redundant peptides were grouped to satisfy the principles of parsimony, and proteins were filtered at a threshold of two peptides per protein and with an FDR of 1%.

#### Batch correction and Outlier Removal

One control peptide that was missing entirely from one of the batches was excluded from further analyses. The initial set of 4,902 protein identifications was then consolidated into 4,034 groups by summing abundances based on the protein names assigned by Scaffold DIA. Missing values were imputed using a minimum value imputation algorithm, and protein abundance values were subsequently log2-transformed.

To correct for technical variation, the RUV4 algorithm [92] (from the R package ruv) was applied using 13 control peptides as negative controls. Following adjustment, proteins with greater than 20% missing values in the original dataset were removed, leaving 1,305 proteins for further analysis. These protein abundance values were then scaled (0 mean, 1 SD) to facilitate comparisons of effect sizes in downstream statistical testing.

Quality control measures included principal component analysis, which identified two subjects as outliers; these subjects were removed from further analyses. The final dataset comprised proteomic data from subjects with the following APOE genotype distribution: APOEe3/3 (n=92), APOEe3/4 (n=193), and APOEe4/4 (n=17).

Neuropathological classification based on dichotomized NIA-Reagan criteria resulted in 81 subjects classified as “No AD” and 221 subjects as “AD.” The mean age at death was 89.8 years, with a gender distribution of 200 females and 102 males.

### DNA Methylation Data Acquisition and Processing

DNA methylation (DNAme) profiling was performed using the Illumina EPIC array, as previously described [93]. To minimize confounding by sex-specific effects, CpG sites located on sex chromosomes, as indicated in the Illumina EPIC manifest, were excluded from further analysis. Furthermore, CpGs in the lowest 10% of variance were removed prior to conducting differential methylation analyses, resulting in a final dataset of 761,813 CpG sites.

The APOE genotype distribution for this cohort was as follows: APOEe3/3 (n=96), APOEe3/4 (n=197), and APOEe4/4 (n=17) and generated as reported [37].

Neuropathological classification based on the dichotomized NIA-Reagan criteria yielded 85 subjects classified as “No AD” and 225 as “AD.” The mean age of the subjects was 89.35 years, with a gender distribution of 205 females and 105 males.

### Genotyping and Polygenic Risk Score (PRS) Calculation

To quantify genetic risk for AD, we derived polygenic risk score (PRS) weights using genome-wide association study (GWAS) summary statistics from Wightman et al. [40]. The GWAS included 39,918 AD cases and 358,140 controls. PRS weights were applied to imputed genome-wide genotypes from 1,780 ROSMAP participants to generate PRS scores, with genotype data accessed from https://pmc.ncbi.nlm.nih.gov/articles/PMC6080491/ [94].

Two versions of PRS were constructed: one excluding genetic variants within the APOE genomic region (GRCh37:19:40,000,000-50,000,000; non-APOE PRS) and another including genome-wide genetic variants (APOE PRS). PRS was calculated using two Bayesian-based methods, PRS-CS-auto and SDPR, both of which do not require parameter tuning [54,55]. The number of genetic variants included in PRS estimation varied by method and APOE inclusion criteria: for PRS-CS-auto, 1,048,147 variants were included in the APOE PRS and 1,044,866 in the non-APOE PRS, while for SDPR, the corresponding numbers were 958,326 and 955,300.

For these analyses, a subset of 254 subjects was used. In this subset, the APOE genotype distribution was as follows: 79 subjects with APOEe3/3, 164 with APOEe3/4, and 11 with APOEe4/4. Neuropathological classification based on the NIA-Reagan score yielded 67 subjects classified as no AD and 187 as AD, with a mean age of 89.70 years (165 females and 89 males).

### Estimation of Neuron Proportion

We used the DNAme data to estimate neuronal proportions in brain tissue samples via the Cell Epigenotype Specific Model (CETS) [101] implemented in R.

### Statistical Analyses

Differential methylation and protein abundance analyses were performed using the limma package [102] (functions lmFit and eBayes), with models adjusted for age, sex, estimated neuronal proportion, and post-mortem interval (PMI). Functional enrichment analysis for differentially abundant proteins was conducted using g:Profiler [103] with false discovery rate (FDR) correction set at 0.05.

Interaction analyses were carried out using logistic regression models of the form: AD ∼ APOE genotype × Protein/DNAme + sex + age + neuronal proportion + PMI implemented with the R stats package. APOE genotype was dichotomized based on the presence of at least one e4 allele.

Comparisons of protein and DNAme levels across subject groups stratified by APOE genotype and AD pathology were performed on the covariate-adjusted molecule values (i.e., residuals from linear regression models, e.g., Protein ∼ sex + age + PMI + neuronal proportion) using the Wilcoxon test.

Mediation analyses were conducted using the R mediation package [104].

### Network Analysis

Protein-protein interaction networks were constructed using the STRING database [52] to elucidate relationships among selected proteins. The CpG site cg06329447 was mapped to the corresponding gene, ELAVL4, for network inclusion. Clustering of the network was performed using the Markov Cluster Algorithm (MCL) [53] with an inflation parameter of 2.4 to identify functional modules within the network.

### Predictive modeling

To evaluate the predictive performance of our molecular features for AD status, we constructed logistic regression models with the following form: AD ∼ Predictors + age + sex where “predictors” represent the candidate molecular features (e.g., proteins, CpG module eigengenes, or PRS scores). To facilitate direct comparison across different omic types, we restricted the analysis to the subset of subjects (n=247) with complete data for proteomics, DNAme, and PRS. In this subset, the APOE genotype distribution was as follows: 76 subjects with APOEe3/3, 160 with APOEe3/4, 11 and 11 with APOEe4/4, with a dichotomized NIA-Reagan classification of 64 “No AD” and 183 “AD” subjects, a mean age of 89.80 years, 161 females, and 86 males.

Using the caret package [105], we performed five-fold cross-validation repeated five times, with out-of-fold predictions for every sample. This approach ensures that every sample contributes to both training and testing in an unbiased manner, and yields stable, smoothed estimates of predictive performance stratified by APOEe4 status. We then extracted false positive and true positive rates and calculated the area under the curve for plotting.

### WGCNA for DNA Methylation

Weighted Gene Co-Expression Network Analysis (WGCNA) [56] was performed on DNAme data to identify modules of co-methylated CpG sites. CpGs were pre-selected based on their association with AD in linear models adjusted for age, sex, estimated neuronal proportion, and PMI, retaining sites with a nominal unadjusted p-value <0.05; a total of 67,654 CpG sites were included in the analysis. Network construction was conducted with the following parameters in the blockwiseModules function: a soft-thresholding power of 6, a signed network type, a minimum module size of 30, and a maximum block size of 34,000. Module eigengenes (defined as the first principal component of each module) were computed to summarize the methylation profiles of each module. Associations between module eigengenes and AD neuropathology were assessed using linear regression models adjusted for age, sex, neuronal proportion, and PMI. Additionally, moderation analyses were performed using logistic regression models structured as: AD ∼ APOE genotype × Eigengene + covariates to evaluate whether the relationship between DNAme modules and AD differed by APOE genotype.

## Data Availability

All generated data and associated metadata have been made publicly available via Synapse.

## ACKNOWLEDGEMENT & FUNDING

This study was funded by the NIA (R01AG057912 to Higgins-Chen, Levine). We also thank the Keck MS & Proteomics Resource at Yale School of Medicine for providing the necessary mass spectrometers and the accompany biotechnology tools funded in part by the Yale School of Medicine and by the Office of The Director, National Institutes of Health (S10OD02365101A1, S10OD019967, and S10OD018034). The funders had no role in study design, data collection and analysis, decision to publish, or preparation of the manuscript. TTL had efforts under the R01AG057912 (Levine, PI). ROSMAP is supported by P30AG10161, P30AG72975, R01AG17917. R01AG015819, U01AG072572, and U01AG046152.

## AUTHOR CONTRIBUTIONS

YM, MEL, AHC conceived and designed the analyses. YM processed and analyzed the data, and wrote the initial manuscript. AHC provided supervision. AP assisted with data analysis. LX accessed whole-genome sequencing data and calculated polygenic risk scores (PRS). WW contributed to generating proteomic data and documentation. KTE accessed, cleaned, and harmonized DNA methylation and phenotypic data. JG, DB, JK, RS, GZ, JF contributed intellectually through manuscript editing and reviewing biological findings. BCC advised on proteomics data processing. HZ provided intellectual input on statistical analyses and project direction. SH, CHvD, CG, DAB aided in the design of the analysis. CHvD, DAB also provided original samples and metadata. TL is a subaward co-investigator and generated proteomic data and documentation. All authors reviewed and edited the manuscript.

