## Supplementary Data for "Beyond the Genotype: A Multi-Omic Analysis of APOEe4’s Role in Alzheimer’s Disease"

### **Supplementary Figure 1**

**Overlap of Subjects Across Proteomics, DNA Methylation, and Polygenic Risk Score Datasets**

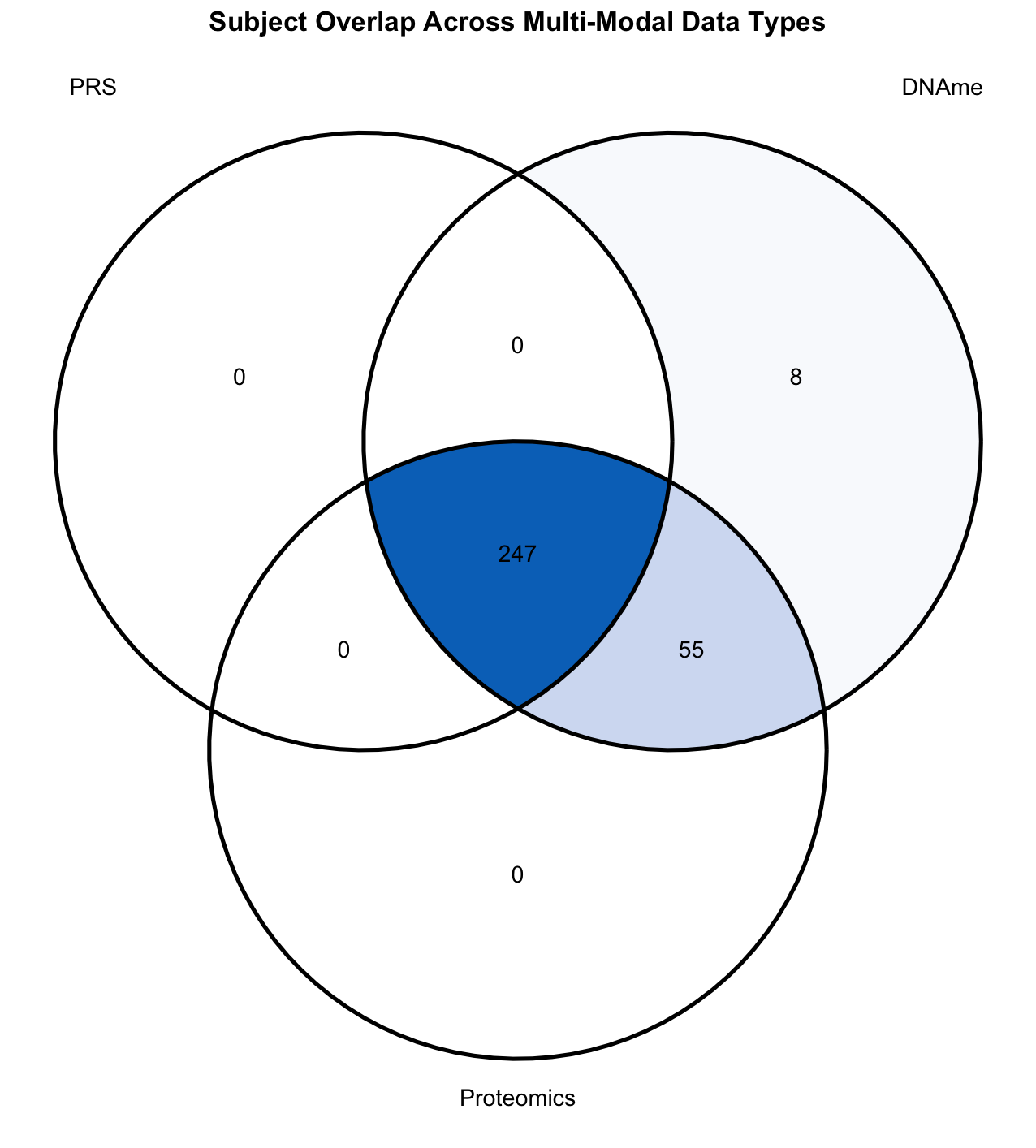

*A venn diagram indicating the overlap of subjects across the three molecular data types used in the study: proteomics, DNA methylation (DNAme), and polygenic risk scores (PRS). Each circle represents the set of individuals for whom that specific data type was available. The intersections indicate subjects with data available in two or more modalities, highlighting the subset of individuals with multi-omic integration potential.*

### **Supplementary Figure 2**

**Gene Ontology (GO) enrichment analysis of proteins differentially abundant in AD versus non-AD samples.**

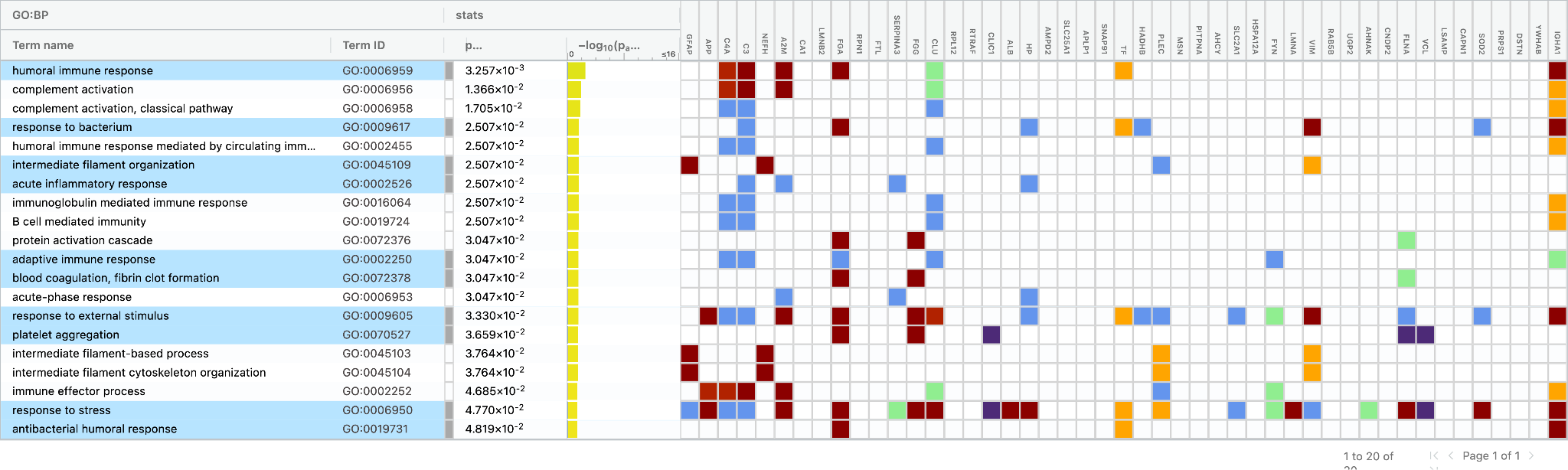

*Results are presented separately for upregulated (top panel) and downregulated (bottom panel) proteins, identified in differential abundance analyses adjusted for age, sex, neuron proportion, and post-mortem interval. GO term enrichment was performed using g:Profiler, and p-values represent adjusted values (multiple testing corrected using FDR). Highlighted in blue are driver terms, identified through g:Profiler's algorithm. Node colors represent the type of evidence supporting the assignment of each protein to the enriched GO term (e.g., experimental, computational), as defined in g:Profiler's evidence code legend.*

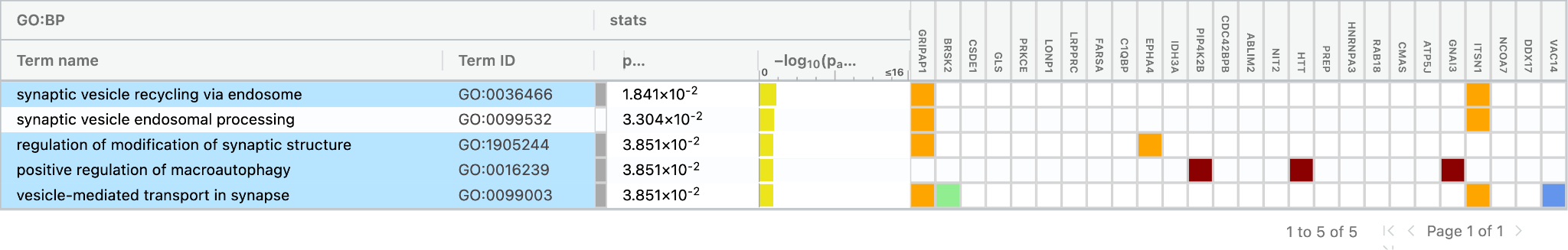

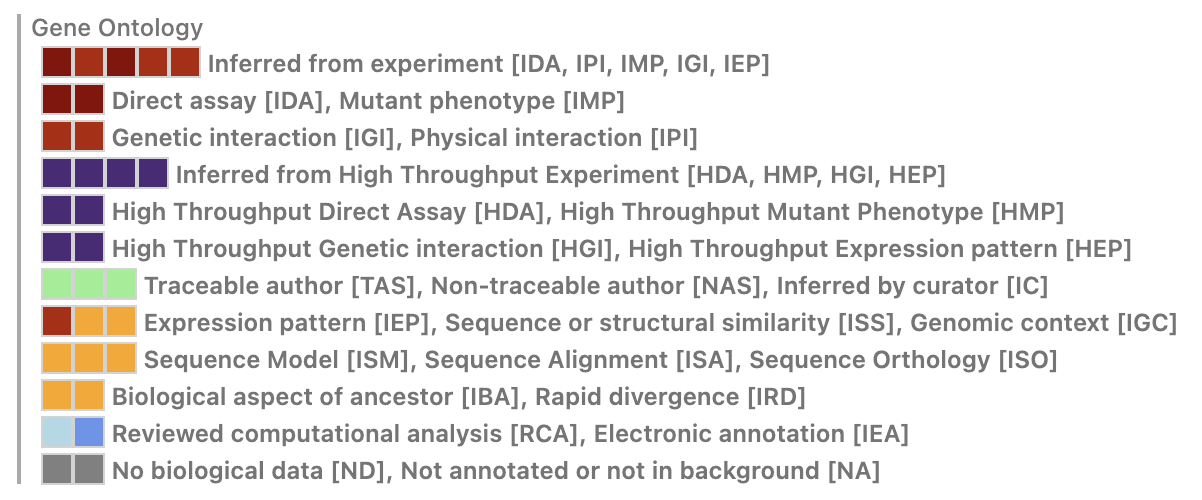

### **Supplementary Table 1**

**Differentially Expressed Proteins in Alzheimer's Disease (AD) vs. Non-AD Samples (upregulated).**

| **Protein** | **logFC** | **P.Value** | **adj.P.Val** |
| --- | --- | --- | --- |
| **GFAP--GFAP** | 1.09701861010002 | 1.86536218952625E-12 | 2.43243229514223E-09 |
| **A4--APP** | 2.52487045343808 | 8.19579655226578E-12 | 5.34365935207729E-09 |
| **CO4A--C4A** | 1.04556222268234 | 8.68210739826775E-09 | 3.77382268244705E-06 |
| **CO3--C3** | 0.92925849670782 | 1.76343373045383E-07 | 5.74879396127948E-05 |
| **NFH--NEFH** | 0.79506694790247 | 1.09097364665185E-06 | 0.000284525927046802 |
| **A2MG--A2M** | 0.715814030295372 | 2.84533175627013E-06 | 0.000618385435029374 |
| **CAH1--CA1** | 0.914037676396953 | 8.68663503454218E-06 | 0.00141592151063037 |
| **LMNB2--LMNB2** | 0.598834057521592 | 1.88612966137462E-05 | 0.0026239921279902 |
| **FIBA--FGA** | 1.00690857118697 | 4.02157046008968E-05 | 0.00413295078251815 |
| **RPN1--RPN1** | 0.751613898227008 | 4.04648828698107E-05 | 0.00413295078251815 |
| **FRIL--FTL** | 0.85423600542123 | 5.01783474887948E-05 | 0.00466527445605807 |
| **AACT--SERPINA3** | 0.877996977122507 | 5.36649669025085E-05 | 0.00466527445605807 |
| **FIBG--FGG** | 0.809423641177068 | 0.000201901798746592 | 0.0148802376465479 |
| **CLUS--CLU** | 0.617561545401129 | 0.00020738326300243 | 0.0148802376465479 |
| **RL12--RPL12** | 0.799757881408709 | 0.000227009052571125 | 0.0148802376465479 |
| **RTRAF--RTRAF** | 0.835023560887372 | 0.00023963572897048 | 0.0148802376465479 |
| **CLIC1--CLIC1** | 0.962987366079608 | 0.000311484985415728 | 0.0164024855024562 |
| **ALBU--ALB** | 0.425631787987853 | 0.000314464829418254 | 0.0164024855024562 |
| **HPT--HP** | 0.723301901808403 | 0.000422501628017055 | 0.0210480880945722 |
| **AMPD2--AMPD2** | 0.636925999699188 | 0.000730470735791089 | 0.031751127982386 |
| **TXTP--SLC25A1** | 0.612965937260826 | 0.000771225273527773 | 0.0324412179574263 |
| **APLP1--APLP1** | 0.948889098239546 | 0.000796657803287645 | 0.0324638054839715 |
| **AP180--SNAP91** | 0.51721054945398 | 0.000927454795946903 | 0.0355372872878407 |
| **TRFE--TF** | 0.437965477875534 | 0.000953838232419037 | 0.0355372872878407 |
| **ECHB--HADHB** | 0.375172927742654 | 0.00105874825873169 | 0.0373137224158413 |
| **PLEC--PLEC** | 0.314842378678693 | 0.00115766307731009 | 0.0397261224424303 |
| **TAU--MAPT** | 0.229964937270226 | 0.001199793668747 | 0.040155754460639 |
| **MOES--MSN** | 0.442229733691021 | 0.00124192426018425 | 0.0405853864788471 |
| **PIPNA--PITPNA** | 0.572521860875135 | 0.00126792735356918 | 0.0405853864788471 |
| **SAHH--AHCY** | 0.636019428680314 | 0.00133832179339756 | 0.0405853864788471 |
| **GTR1--SLC2A1** | 0.62671159839351 | 0.00146261189515764 | 0.0414618676366426 |
| **A0A1B0GTF3--HSPA12A** | 0.452975405721789 | 0.00160334158859364 | 0.0444842006707683 |
| **FYN--FYN** | 0.72226466856335 | 0.00188921989312129 | 0.0503982065294256 |
| **LMNA--LMNA** | 0.513199369107292 | 0.00189379763799222 | 0.0503982065294256 |
| **VIME--VIM** | 0.381908305783373 | 0.00195505722780881 | 0.0509878925012538 |
| **H31T--HIST3H3** | 0.318073762822049 | 0.00209820977506317 | 0.0536483440525956 |
| **RAB5B--RAB5B** | 0.426775867013343 | 0.00231635525617726 | 0.0565265447261682 |
| **UGPA--UGP2** | 0.485542879985982 | 0.00234082317117568 | 0.0565265447261682 |
| **AHNK--AHNAK** | 0.405900234951988 | 0.00260439182775903 | 0.060645123989246 |
| **CNDP2--CNDP2** | 0.389702921231243 | 0.00265563758482165 | 0.0607535335194287 |
| **FLNA--FLNA** | 0.473324539258714 | 0.00296726654544619 | 0.0667123375045143 |
| **VINC--VCL** | 0.355496788803123 | 0.00349192228431666 | 0.0771774009957445 |
| **LSAMP--LSAMP** | 0.603792312946749 | 0.00373024621830024 | 0.0774897254238891 |
| **CAN1--CAPN1** | 0.427317954392241 | 0.00404563823064146 | 0.0799320038296434 |
| **SODM--SOD2** | 0.47627447413486 | 0.00412352585680275 | 0.080254891302549 |
| **PRPS1--PRPS1** | 0.597716767494925 | 0.00500405296853338 | 0.0907122597026781 |
| **DEST--DSTN** | 0.822280475553355 | 0.00500865237622149 | 0.0907122597026781 |
| **1433B--YWHAB** | 0.675952871465861 | 0.00518500918120726 | 0.0913682698958684 |
| **IGHA1--IGHA1** | 0.556742484401045 | 0.00544998789482301 | 0.0947571228646561 |

*These tables list proteins with significant changes in abundance identified in the proteomics dataset. Differential expression was assessed using linear models adjusted for age, sex, neuron proportion, and post-mortem interval. p-values were corrected for multiple testing using the false discovery rate (FDR) method. Protein refers to the UniProt identifier and corresponding gene symbol. logFC denotes the log₂ fold change in protein abundance between AD and non-AD groups. The top table presents proteins upregulated in AD; the bottom table includes downregulated proteins. adj.P.Val indicates FDR-adjusted p-values.*

**Differentially Expressed Proteins in Alzheimer's Disease (AD) vs. Non-AD Samples (downregulated).**

| **Protein** | **logFC** | **P.Value** | **adj.P.Val** |
| --- | --- | --- | --- |
| **GRAP1--GRIPAP1** | -1.36570910710302 | 6.36307511133198E-06 | 0.00118534999216813 |
| **BRSK2--BRSK2** | -1.55013069267292 | 2.01226390183297E-05 | 0.0026239921279902 |
| **CSDE1--CSDE1** | -0.984069510318382 | 4.12027301938159E-05 | 0.00413295078251815 |
| **GLSK--GLS** | -0.642471095412722 | 0.000118070427196502 | 0.00962273981651494 |
| **KPCE--PRKCE** | -0.966285127781934 | 0.000234156009903957 | 0.0148802376465479 |
| **LONM--LONP1** | -0.637695689116267 | 0.000267442777901319 | 0.0153706591503702 |
| **LPPRC--LRPPRC** | -0.678262418970709 | 0.000271108251885364 | 0.0153706591503702 |
| **SYFA--FARSA** | -0.658110566422651 | 0.000435811639994976 | 0.0210480880945722 |
| **C1QBP--C1QBP** | -0.776668030540852 | 0.000571427826889177 | 0.026612210223696 |
| **EPHA4--EPHA4** | -0.653508489651452 | 0.000671172248268572 | 0.0301796073014558 |
| **IDH3A--IDH3A** | -0.537309381464288 | 0.000916211215642278 | 0.0355372872878407 |
| **PI42B--PIP4K2B** | -0.709601691965666 | 0.00104593792080408 | 0.0373137224158413 |
| **MRCKB--CDC42BPB** | -0.805137522762012 | 0.0013215585803977 | 0.0405853864788471 |
| **ABLM2--ABLIM2** | -0.606009388197029 | 0.00133545094473415 | 0.0405853864788471 |
| **NIT2--NIT2** | -0.752881185824413 | 0.00138335014605254 | 0.0409974679648299 |
| **HD--HTT** | -0.740252704156867 | 0.00145361814337824 | 0.0414618676366426 |
| **PPCE--PREP** | -0.609186900626872 | 0.00218188527885516 | 0.0547149693005218 |
| **ROA3--HNRNPA3** | -0.567211828318682 | 0.00251916646145165 | 0.05972714664969 |
| **RAB18--RAB18** | -0.469047866898366 | 0.00370080185476467 | 0.0774897254238891 |
| **NEUA--CMAS** | -0.660490503382414 | 0.00370975684744423 | 0.0774897254238891 |
| **ATP5J--ATP5J** | -0.752686412724627 | 0.00374375207185967 | 0.0774897254238891 |
| **GNAI3--GNAI3** | -0.572687431387441 | 0.00389622245942414 | 0.0784453041839314 |
| **ITSN1--ITSN1** | -0.741299941703172 | 0.0039102337208248 | 0.0784453041839314 |
| **NCOA7--NCOA7** | -0.795620354324531 | 0.00450614339648778 | 0.0855132982005829 |
| **USMG5--USMG5** | -0.495786377261206 | 0.00452486010417195 | 0.0855132982005829 |
| **DDX17--DDX17** | -0.536543255945199 | 0.00490643117770033 | 0.0907122597026781 |
| **VAC14--VAC14** | -0.747748639081647 | 0.00508393395138333 | 0.0908143818164912 |

### **Supplementary Table 2**

**Differentially methylated CpG sites in Alzheimer's disease (AD) vs. non-AD samples.**

| **chr** | **pos** | **strand** | **Name** | **Gencode_Group** | **ΔBeta** | **AveExpr** | **P.Value** | **adj.P.Val** | **Gene** | **APOE_Interaction_Significance** |
| --- | --- | --- | --- | --- | --- | --- | --- | --- | --- | --- |
| chr1 | 167090618 | - | cg04138112 |  | -0.0196644167945514 | 0.0638331601497997 | 8.67801182828472E-09 | 0.00661102222494107 |  | 0.340148381199636 |
| chr8 | 41519308 | - | cg05066959 | ExonBnd | 0.0295239536332871 | 0.907085253814065 | 4.42044586241717E-08 | 0.016837765618928 | ANK1 | 0.210302106013635 |
| chr17 | 74475402 | - | cg12309456 |  | 0.0113664138298963 | 0.965699429081036 | 1.9516922111857E-07 | 0.0454063317643492 |  | 0.792177319584915 |
| chr11 | 69464012 | + | cg15974867 |  | 0.013917609715574 | 0.915521029068693 | 2.38411955502724E-07 | 0.0454063317643492 | CCND1 | 0.879073920212815 |
| chr6 | 41103849 | + | cg11915672 | 5'UTR | 0.0126736499133197 | 0.805014661293109 | 4.04603271360778E-07 | 0.0462677343016853 |  | 0.292568839239632 |
| chr10 | 22623047 | - | cg03760191 |  | 0.0221472205783922 | 0.810698195983497 | 4.26291982689832E-07 | 0.0462677343016853 |  | 0.259327920981804 |
| chr19 | 49962324 | - | cg20618448 | 5'UTR;ExonBnd | 0.00609003380014371 | 0.974655124090867 | 4.74558485032848E-07 | 0.0462677343016853 | ALDH16A1 | 0.720214325147132 |
| chr1 | 167090646 | - | cg17104258 |  | -0.0254407865603755 | 0.0541795252012572 | 4.85869727102953E-07 | 0.0462677343016853 |  | 0.716923192509315 |
| chr6 | 6611038 | + | cg06701353 | 3'UTR | 0.00701410699638838 | 0.94085876520665 | 7.37340232549781E-07 | 0.0624128193977163 | LY86-AS1 | 0.686078017263644 |
| chr19 | 47249203 | + | cg04247584 | 5'UTR;5'UTR;TSS200 | -0.00540609537801536 | 0.0426945070930958 | 8.8043960051424E-07 | 0.0670730333386555 | STRN4 | 0.998554669861611 |
| chr1 | 43296491 | - | cg25285237 | 5'UTR;1stExon;5'UTR | 0.0174538310910972 | 0.854768353400614 | 1.2617443814216E-06 | 0.0829826148218833 | ERMAP | 0.138049461770083 |
| chr1 | 43296526 | + | cg12410370 | 5'UTR;1stExon;5'UTR | 0.00864006428093644 | 0.94365532783575 | 1.30713361134898E-06 | 0.0829826148218833 | ERMAP | 0.0985613963721393 |
| chr2 | 123819265 | - | cg20178603 |  | 0.0132702688095822 | 0.882952134361383 | 1.44675422724966E-06 | 0.0832372445277485 |  | 0.339284351371934 |
| chr2 | 242553049 | - | cg25230945 |  | 0.0321057318685144 | 0.491501647087621 | 1.52966859765911E-06 | 0.0832372445277485 |  | 0.627261086709505 |
| chr5 | 175975901 | + | cg00332146 | TSS1500;5'UTR | -0.0205611709872232 | 0.656218500452585 | 2.26789332320871E-06 | 0.099987030766679 | CDHR2 | 0.155709333643554 |
| chr20 | 42762143 | + | cg20134175 |  | 0.0172207481274379 | 0.861966421738308 | 2.68985338448591E-06 | 0.099987030766679 |  | 0.523584336433327 |
| chr2 | 233963033 | - | cg10609370 |  | 0.00937098083594856 | 0.921151724030391 | 2.81116938856187E-06 | 0.099987030766679 | INPP5D | 0.284840112528735 |
| chr17 | 74475270 | - | cg05810363 |  | 0.0141462119037574 | 0.968383524718205 | 2.97588318882248E-06 | 0.099987030766679 |  | 0.730413534258425 |
| chr17 | 74475355 | + | cg12163800 | ExonBnd | 0.0137052622067001 | 0.94690758524788 | 3.136345889543E-06 | 0.099987030766679 | RHBDF2 | 0.453084134062784 |
| chr5 | 116040655 | - | cg02870855 |  | 0.00984201872564556 | 0.92021619964367 | 3.34992440957003E-06 | 0.099987030766679 |  | 0.193815010768101 |
| chr7 | 101398184 | + | cg14398883 |  | -0.0104442036077854 | 0.0457602635551518 | 3.42009201304579E-06 | 0.099987030766679 |  | 0.565304930142053 |
| chr19 | 3178955 | - | cg17518965 | 1stExon | -0.0173438046527316 | 0.728183061531577 | 3.5062448704158E-06 | 0.099987030766679 | S1PR4 | 0.189817374772304 |
| chr12 | 72015410 | - | cg26385186 |  | 0.0157266545887734 | 0.822035781665881 | 3.58195424125777E-06 | 0.099987030766679 |  | 0.330036681764709 |
| chr1 | 50574241 | + | cg06329447 | TSS1500;TSS1500;TSS1500;TSS1500 | 0.0131361297459284 | 0.588627904147719 | 3.60244173314095E-06 | 0.099987030766679 | ELAVL4 | 0.0328207267583745 |
| chr17 | 74475294 | + | cg13076843 |  | 0.0171309841212702 | 0.904980981220305 | 3.6093841522355E-06 | 0.099987030766679 |  | 0.934567214563585 |

*This table lists CpG sites with significant differences in DNA methylation between AD and non-AD individuals, identified through linear modeling adjusted for age, sex, neuron proportion, and post-mortem interval. Results are filtered at FDR < 0.10 threshold. chr and pos indicate the chromosome and genomic position of each CpG site. strand refers to the DNA strand. Name is the CpG probe ID. Gencode_Group denotes the annotated genomic feature category (e.g., TSS200, gene body). ΔBeta represents the mean methylation difference (AD – non-AD). AveExpr is the average methylation signal intensity. P.Value and adj.P.Val are the raw and FDR-adjusted p-values, respectively. Gene indicates the gene symbol annotated to the CpG site. APOE_Interaction_Significance notes whether the CpG site also showed a nominally significant interaction with APOEe4 genotype in the moderation analysis.*

### Supplementary Table 3

| ***APOE/AD***  ***status*** | ***N*** | ***Age***  ***Mean (SD)*** | ***Female%*** | ***CERAD status*** | | ***Braak stage*** | | | | ***Molecular data*** | |
| --- | --- | --- | --- | --- | --- | --- | --- | --- | --- | --- | --- |
|  |  |  |  | ***Non-AD*** | ***AD*** | ***0*** | ***I-II*** | ***III-IV*** | ***V-VI*** | ***Prot*** | ***DNAm*** |
| ***e4− / AD−*** | ***43*** | ***87.3 (6.0)*** | ***62.8%*** | ***40*** | ***3*** | ***2*** | ***15*** | ***26*** | ***0*** | ***40*** | ***43*** |
| ***e4− / AD+*** | ***53*** | ***91.2 (4.7)*** | ***60.4%*** | ***0*** | ***53*** | ***0*** | ***1*** | ***30*** | ***22*** | ***52*** | ***53*** |
| ***e4+ / AD−*** | ***43*** | ***88.8 (6.9)*** | ***62.7%*** | ***37*** | ***5*** | ***0*** | ***16*** | ***25*** | ***1*** | ***41*** | ***42*** |
| ***e4+ / AD+*** | ***172*** | ***89.5 (5.7)*** | ***69.8%*** | ***0*** | ***172*** | ***1*** | ***6*** | ***67*** | ***98*** | ***169*** | ***172*** |

### Supplementary Table 4

**Proteins showing significant APOEe4 × Alzheimer's disease (AD) interaction effects.**

| Protein | Beta_Interaction | SE_Interaction | P_Interaction | Adj_P_Interaction |
| --- | --- | --- | --- | --- |
| BSN--BSN | 2.2751780 | 0.6083254 | 0.0001839707 | 0.04904426 |
| CAP2--CAP2 | 1.9142761 | 0.5119829 | 0.0001847940 | 0.04904426 |
| CSKI1--CASKIN1 | 1.8851293 | 0.5110065 | 0.0002250914 | 0.04904426 |
| DHE3--GLUD1 | 2.0292492 | 0.5672846 | 0.0003473909 | 0.04904426 |
| SFXN1--SFXN1 | 1.7635566 | 0.4977589 | 0.0003956118 | 0.04904426 |
| HSP7C--HSPA8 | 1.7913992 | 0.5057259 | 0.0003967540 | 0.04904426 |
| AP2A1--AP2A1 | 1.9990532 | 0.5679729 | 0.0004321526 | 0.04904426 |
| MAON--ME3 | 1.5565084 | 0.4429230 | 0.0004411247 | 0.04904426 |
| AUXI--DNAJC6 | 1.9577033 | 0.5628683 | 0.0005050118 | 0.04904426 |
| LETM1--LETM1 | 1.4882948 | 0.4294098 | 0.0005284441 | 0.04904426 |
| AAK1--AAK1 | 1.9972341 | 0.5787738 | 0.0005589221 | 0.04904426 |
| SYSM--SARS2 | 1.4713969 | 0.4272750 | 0.0005738609 | 0.04904426 |
| HSP74--HSPA4 | 1.6141049 | 0.4687859 | 0.0005749580 | 0.04904426 |
| MAP1A--MAP1A | 1.9949895 | 0.5819526 | 0.0006078297 | 0.04904426 |
| OXR1--OXR1 | 1.5073705 | 0.4444398 | 0.0006948093 | 0.04905607 |
| KIF5C--KIF5C | 1.7918813 | 0.5303974 | 0.0007291557 | 0.04905607 |
| AHNK--AHNAK | 1.3083706 | 0.3881893 | 0.0007504687 | 0.04905607 |
| VP26B--VPS26B | 1.3898063 | 0.4129566 | 0.0007640451 | 0.04905607 |
| PFKAP--PFKP | 1.7550131 | 0.5233380 | 0.0007979686 | 0.04905607 |
| SYFB--FARSB | 1.4659432 | 0.4438281 | 0.0009567256 | 0.05370142 |
| BIP--HSPA5 | 1.5097839 | 0.4604397 | 0.0010417396 | 0.05549716 |
| CH60--HSPD1 | 1.5186120 | 0.4643781 | 0.0010746933 | 0.05549716 |
| GHC1--SLC25A22 | 1.2420231 | 0.3832591 | 0.0011924166 | 0.05920807 |
| PSMD2--PSMD2 | 1.2902345 | 0.4051202 | 0.0014484449 | 0.06436006 |
| SFXN5--SFXN5 | 1.4286338 | 0.4503243 | 0.0015115554 | 0.06436006 |
| AFG32--AFG3L2 | 1.1797080 | 0.3735501 | 0.0015880183 | 0.06436006 |
| HEBP1--HEBP1 | 1.2133248 | 0.3843563 | 0.0015952674 | 0.06436006 |
| NRCAM--NRCAM | 1.4456559 | 0.4581457 | 0.0016025054 | 0.06436006 |
| BIG3--ARFGEF3 | 1.4397942 | 0.4572812 | 0.0016405609 | 0.06436006 |
| VAMP1--VAMP1 | 1.4829960 | 0.4718967 | 0.0016743843 | 0.06436006 |
| BASI--BSG | 1.6486124 | 0.5251956 | 0.0016949976 | 0.06436006 |
| STIP1--STIP1 | 1.2613731 | 0.4040410 | 0.0017969191 | 0.06628065 |
| SYNPO--SYNPO | 1.3404772 | 0.4337460 | 0.0019984283 | 0.07166586 |
| PACN1--PACSIN1 | 1.3098648 | 0.4275065 | 0.0021842482 | 0.07301965 |
| IDH3A--IDH3A | 1.6345895 | 0.5340020 | 0.0022058609 | 0.07301965 |
| HXK1--HK1 | 1.4626070 | 0.4801042 | 0.0023157135 | 0.07394945 |
| NCOA7--NCOA7 | 1.5301062 | 0.5041762 | 0.0024064415 | 0.07394945 |
| VATD--ATP6V1D | 1.2941943 | 0.4275672 | 0.0024709264 | 0.07394945 |
| XPO1--XPO1 | 1.1043788 | 0.3665891 | 0.0025903786 | 0.07394945 |
| NDUS1--NDUFS1 | 1.4192135 | 0.4731468 | 0.0027040467 | 0.07394945 |
| GNAI2--GNAI2 | 1.2443871 | 0.4168214 | 0.0028318903 | 0.07394945 |
| SSBP--SSBP1 | 1.2971127 | 0.4345047 | 0.0028333036 | 0.07394945 |
| CNTN2--CNTN2 | 1.5501475 | 0.5198408 | 0.0028640375 | 0.07394945 |
| NRX3A--NRXN3 | 1.1181428 | 0.3764780 | 0.0029779205 | 0.07454254 |
| USO1--USO1 | 1.2301207 | 0.4145822 | 0.0030059068 | 0.07454254 |
| KAP3--PRKAR2B | 1.3109944 | 0.4426609 | 0.0030602280 | 0.07454254 |
| NCAM2--NCAM2 | 1.4095032 | 0.4774096 | 0.0031531597 | 0.07538387 |
| ARP2--ACTR2 | 1.2451440 | 0.4228751 | 0.0032350582 | 0.07542507 |
| AGK--AGK | 1.2287751 | 0.4178111 | 0.0032717303 | 0.07542507 |
| ACLY--ACLY | 1.2529081 | 0.4293701 | 0.0035226827 | 0.07841006 |
| EF2--EEF2 | 1.3409068 | 0.4616285 | 0.0036756813 | 0.08042889 |
| HPCL4--HPCAL4 | 1.2644634 | 0.4378851 | 0.0038811858 | 0.08251196 |
| RAB4A--RAB4A | 1.2182940 | 0.4224051 | 0.0039242983 | 0.08251196 |
| DMXL2--DMXL2 | 1.1855758 | 0.4134029 | 0.0041327679 | 0.08361344 |
| PI4KA--PI4KA | 1.2353478 | 0.4340924 | 0.0044297574 | 0.08798180 |
| AL7A1--ALDH7A1 | 1.2880134 | 0.4536993 | 0.0045267367 | 0.08854571 |
| PLPHP--PLPBP | 1.3725051 | 0.4877050 | 0.0048897048 | 0.09421804 |
| MIC19--CHCHD3 | 0.9783611 | 0.3498262 | 0.0051626411 | 0.09659376 |
| AT1A1--ATP1A1 | 1.1645783 | 0.4213384 | 0.0057097994 | 0.09969738 |
| VGF--VGF | 1.0586093 | 0.3837485 | 0.0058049257 | 0.09969738 |
| BAIP2--BAIAP2 | 1.3858466 | 0.5038238 | 0.0059475835 | 0.09969738 |
| E41L1--EPB41L1 | 1.1396807 | 0.4144172 | 0.0059580550 | 0.09969738 |
| APEX1--APEX1 | 1.1652196 | 0.4250432 | 0.0061175269 | 0.09969738 |
| GSTK1--GSTK1 | 1.4071239 | 0.5135322 | 0.0061422555 | 0.09969738 |
| CNTN1--CNTN1 | 1.4192221 | 0.5195659 | 0.0063036504 | 0.09969738 |
| PLCB1--PLCB1 | 1.3275120 | 0.4863878 | 0.0063463531 | 0.09969738 |
| SH3G2--SH3GL2 | 1.2551491 | 0.4602832 | 0.0063931168 | 0.09969738 |
| TPIS--TPI1 | 1.0801853 | 0.3966138 | 0.0064590451 | 0.09969738 |
| CRKL--CRKL | 0.9138542 | 0.3360687 | 0.0065430578 | 0.09969738 |
| CRYM--CRYM | 1.1966216 | 0.4402282 | 0.0065641190 | 0.09969738 |
| SYN3--SYN3 | 1.1227709 | 0.4165353 | 0.0070283117 | 0.10203519 |
| WDR1--WDR1 | 1.1256493 | 0.4180500 | 0.0070893292 | 0.10203519 |
| LRC47--LRRC47 | 1.1615276 | 0.4320679 | 0.0071817102 | 0.10203519 |
| ES1--C21orf33 | 0.8849781 | 0.3305400 | 0.0074202429 | 0.10203519 |
| SEPT8--SEPT8 | 1.1686658 | 0.4373487 | 0.0075364583 | 0.10203519 |
| RYR2--RYR2 | 0.9091796 | 0.3405741 | 0.0075953018 | 0.10203519 |
| ABR--ABR | 1.1378240 | 0.4264893 | 0.0076330504 | 0.10203519 |
| GNAO--GNAO1 | 1.0815213 | 0.4056524 | 0.0076730448 | 0.10203519 |
| OLA1--OLA1 | 1.0815251 | 0.4059892 | 0.0077234565 | 0.10203519 |
| KBTBB--KBTBD11 | 1.2025582 | 0.4519363 | 0.0077931674 | 0.10203519 |
| HS105--HSPH1 | 1.2710133 | 0.4779056 | 0.0078245418 | 0.10203519 |
| BAG6--BAG6 | 1.0331969 | 0.3893938 | 0.0079697824 | 0.10260026 |
| S12A5--SLC12A5 | 1.0685696 | 0.4030909 | 0.0080268215 | 0.10260026 |
| GNAI3--GNAI3 | 1.1381086 | 0.4298997 | 0.0081117750 | 0.10266962 |
| ACPH--APEH | 1.1491124 | 0.4354647 | 0.0083195492 | 0.10293137 |
| SPTN2--SPTBN2 | 1.1837061 | 0.4489266 | 0.0083705142 | 0.10293137 |
| IF4B--EIF4B | 1.0106295 | 0.3837970 | 0.0084574590 | 0.10293137 |
| CADH2--CDH2 | 1.0476584 | 0.3983620 | 0.0085406077 | 0.10293137 |
| CNTP1--CNTNAP1 | 1.2283520 | 0.4673415 | 0.0085792064 | 0.10293137 |
| HUWE1--HUWE1 | 1.0714912 | 0.4090707 | 0.0088102729 | 0.10293137 |
| PGCB--BCAN | 1.3089036 | 0.5011399 | 0.0090053044 | 0.10293137 |
| ANK3--ANK3 | 1.1789164 | 0.4523265 | 0.0091515733 | 0.10293137 |
| STXB5--STXBP5 | 1.0768714 | 0.4135303 | 0.0092117776 | 0.10293137 |
| NP1L1--NAP1L1 | 1.0805424 | 0.4152082 | 0.0092570744 | 0.10293137 |
| ARPC4--ARPC4 | 0.9373443 | 0.3608602 | 0.0093897596 | 0.10293137 |
| AT2B2--ATP2B2 | 1.0970881 | 0.4223599 | 0.0093899859 | 0.10293137 |
| GRM2--GRM2 | 1.2252438 | 0.4723842 | 0.0094936988 | 0.10293137 |
| GIT1--GIT1 | 1.1487171 | 0.4434531 | 0.0095866849 | 0.10293137 |
| FAS--FASN | 1.2175208 | 0.4708687 | 0.0097184101 | 0.10293137 |
| SEPT9--SEPT9 | 0.9935300 | 0.3847689 | 0.0098187757 | 0.10293137 |
| GNL1--GNL1 | 1.2526123 | 0.4851084 | 0.0098192985 | 0.10293137 |
| NDUS2--NDUFS2 | 1.0876874 | 0.4213452 | 0.0098382296 | 0.10293137 |
| FPPS--FDPS | 0.9858117 | 0.3820758 | 0.0098758315 | 0.10293137 |
| FIBG--FGG | 0.8525662 | 0.3304810 | 0.0098865145 | 0.10293137 |
| TPPP--TPPP | 1.0472554 | 0.4068644 | 0.0100539975 | 0.10312310 |
| ACON--ACO2 | 1.2582185 | 0.4888945 | 0.0100646872 | 0.10312310 |
| ALDOC--ALDOC | 1.0997577 | 0.4280134 | 0.0101861040 | 0.10320596 |
| EPHA4--EPHA4 | 1.1037944 | 0.4302253 | 0.0102991836 | 0.10320596 |
| IDHC--IDH1 | 0.8985528 | 0.3502901 | 0.0103126015 | 0.10320596 |
| PSMD1--PSMD1 | 0.8848598 | 0.3459725 | 0.0105396880 | 0.10466721 |
| HPRT--HPRT1 | 1.0331409 | 0.4045490 | 0.0106551647 | 0.10500624 |
| LRP1--LRP1 | 1.0803117 | 0.4239492 | 0.0108277141 | 0.10589832 |
| GUAD--GDA | 1.0281487 | 0.4048693 | 0.0111024310 | 0.10776871 |
| AMPH--AMPH | 1.0085111 | 0.3979827 | 0.0112750245 | 0.10858859 |
| NAKD2--NADK2 | 0.8519705 | 0.3366751 | 0.0113886469 | 0.10858859 |
| DPP3--DPP3 | 1.1992699 | 0.4746803 | 0.0115212234 | 0.10858859 |
| KI21A--KIF21A | 1.0690840 | 0.4237543 | 0.0116395889 | 0.10858859 |
| ODO2--DLST | 1.0504793 | 0.4167373 | 0.0117113976 | 0.10858859 |
| CLH1--CLTC | 1.1924518 | 0.4732279 | 0.0117412988 | 0.10858859 |
| CK054--C11orf54 | 0.9428620 | 0.3747034 | 0.0118597916 | 0.10858859 |
| GRAP1--GRIPAP1 | 1.3148591 | 0.5238648 | 0.0120758260 | 0.10978797 |

*This table lists proteins that exhibited significant interaction effects between APOEe4 genotype and AD status in the proteomics dataset. Interaction effects were estimated using linear models adjusted for sex, age, neuron proportion, post-mortem interval. Protein refers to the UniProt identifier and gene symbol. Beta_Interaction represents the estimated interaction effect size. SE_Interaction is the standard error of the interaction coefficient. P_Interaction and Adj_P_Interaction are the nominal and FDR-adjusted p-values, respectively. The full list includes all proteins with interaction effects at FDR < 0.10 to facilitate biological interpretation and network integration.*

### **Supplementary** **Table 5**

**Mediation analysis of APOEe4's effect on Alzheimer's disease (AD) through protein abundance.**

| Protein | ACME | ACME_p | Prop_Mediated | ADE | ADE_p | Total_Effect |
| --- | --- | --- | --- | --- | --- | --- |
| GRAP1--GRIPAP1 | 0.0790846842 | 0.000 | 0.340165797 | 0.1534040 | 0.004 | 0.2324886 |
| VAMP1--VAMP1 | 0.0373385395 | 0.002 | 0.147893246 | 0.2151310 | 0.000 | 0.2524695 |
| DPP3--DPP3 | 0.0317052601 | 0.004 | 0.126000962 | 0.2199219 | 0.000 | 0.2516271 |
| CSKI1--CASKIN1 | 0.0382922507 | 0.006 | 0.168173572 | 0.1894026 | 0.000 | 0.2276948 |
| GSTK1--GSTK1 | 0.0637056838 | 0.006 | 0.258406016 | 0.1828276 | 0.002 | 0.2465333 |
| FIBG--FGG | 0.0228655697 | 0.032 | 0.101689064 | 0.2019921 | 0.000 | 0.2248577 |
| SYN3--SYN3 | 0.0226794263 | 0.032 | 0.088755271 | 0.2328482 | 0.000 | 0.2555277 |
| GRM2--GRM2 | 0.0308481472 | 0.064 | 0.130455263 | 0.2056172 | 0.000 | 0.2364653 |
| SYSM--SARS2 | 0.0171549457 | 0.096 | 0.067074848 | 0.2386033 | 0.000 | 0.2557582 |
| BSN--BSN | 0.0246159546 | 0.110 | 0.114804498 | 0.1898003 | 0.000 | 0.2144163 |
| GNAI2--GNAI2 | 0.0103377988 | 0.144 | 0.044045755 | 0.2243681 | 0.000 | 0.2347059 |
| BIG3--ARFGEF3 | 0.0181875100 | 0.146 | 0.078902559 | 0.2123184 | 0.000 | 0.2305060 |
| SYFB--FARSB | 0.0164081407 | 0.168 | 0.073538340 | 0.2067155 | 0.000 | 0.2231236 |
| HSP7C--HSPA8 | -0.0153470502 | 0.196 | -0.066043274 | 0.2477258 | 0.000 | 0.2323787 |
| PI4KA--PI4KA | 0.0132065032 | 0.238 | 0.053612229 | 0.2331273 | 0.000 | 0.2463338 |
| LRC47--LRRC47 | -0.0125857079 | 0.242 | -0.056391622 | 0.2357697 | 0.000 | 0.2231840 |
| MAON--ME3 | 0.0105852383 | 0.314 | 0.045772273 | 0.2206735 | 0.000 | 0.2312587 |
| S12A5--SLC12A5 | -0.0117696624 | 0.338 | -0.048922427 | 0.2523477 | 0.000 | 0.2405781 |
| SFXN1--SFXN1 | -0.0108331881 | 0.400 | -0.047496087 | 0.2389191 | 0.000 | 0.2280859 |
| CAP2--CAP2 | -0.0128139332 | 0.404 | -0.061589790 | 0.2208668 | 0.000 | 0.2080529 |
| RAB4A--RAB4A | -0.0096003880 | 0.416 | -0.040703051 | 0.2454645 | 0.000 | 0.2358641 |
| LRP1--LRP1 | -0.0051675646 | 0.434 | -0.019930995 | 0.2644403 | 0.000 | 0.2592728 |
| DMXL2--DMXL2 | 0.0076072855 | 0.466 | 0.034105941 | 0.2154414 | 0.000 | 0.2230487 |
| GNAO--GNAO1 | -0.0143535569 | 0.476 | -0.056114714 | 0.2701431 | 0.000 | 0.2557895 |
| OXR1--OXR1 | 0.0094814591 | 0.478 | 0.043461639 | 0.2086755 | 0.000 | 0.2181570 |
| ODO2--DLST | 0.0062440130 | 0.528 | 0.027874314 | 0.2177620 | 0.000 | 0.2240060 |
| VP26B--VPS26B | 0.0042774059 | 0.528 | 0.018386148 | 0.2283654 | 0.000 | 0.2326428 |
| NRX3A--NRXN3 | 0.0043801384 | 0.544 | 0.018108460 | 0.2375034 | 0.000 | 0.2418835 |
| HEBP1--HEBP1 | -0.0040801976 | 0.552 | -0.017259099 | 0.2404887 | 0.000 | 0.2364085 |
| KIF5C--KIF5C | 0.0077055896 | 0.606 | 0.034851671 | 0.2133911 | 0.000 | 0.2210967 |
| APEX1--APEX1 | -0.0056209521 | 0.620 | -0.022472913 | 0.2557422 | 0.000 | 0.2501212 |
| SYNPO--SYNPO | 0.0045399121 | 0.642 | 0.020560402 | 0.2162686 | 0.000 | 0.2208085 |
| HPRT--HPRT1 | 0.0037607982 | 0.666 | 0.015483912 | 0.2391234 | 0.000 | 0.2428842 |
| AHNK--AHNAK | 0.0035489468 | 0.680 | 0.013685292 | 0.2557767 | 0.000 | 0.2593256 |
| CRKL--CRKL | 0.0028470078 | 0.712 | 0.011744085 | 0.2395736 | 0.000 | 0.2424206 |
| VGF--VGF | 0.0023978693 | 0.740 | 0.009828404 | 0.2415756 | 0.000 | 0.2439734 |
| GHC1--SLC25A22 | 0.0023722119 | 0.748 | 0.009976034 | 0.2354189 | 0.000 | 0.2377911 |
| BAG6--BAG6 | -0.0007138173 | 0.818 | -0.002832901 | 0.2526878 | 0.000 | 0.2519739 |
| DHE3--GLUD1 | 0.0008479086 | 0.984 | 0.003449832 | 0.2449347 | 0.000 | 0.2457826 |

*This table presents results from mediation analyses assessing whether specific proteins mediate the relationship between APOEe4 genotype and AD status. Analyses were conducted using linear models adjusted for age, sex, neuron proportion, and post-mortem interval. Protein refers to the UniProt identifier and gene symbol. ACME (Average Causal Mediation Effect) estimates the indirect effect of APOEe4 on AD through the protein. ACME_p is the p-value for the mediation effect. Prop_Mediated indicates the proportion of the total effect mediated by the protein. ADE (Average Direct Effect) represents the direct effect of APOEe4 on AD not explained by the protein, with ADE_p as its p-value. Total_Effect reflects the sum of direct and mediated effects. Proteins with ACME_p < 0.05 were considered significant mediators.*

### **Supplementary Table 6**

**Associations between prioritized proteins and neuropathological burden (CERAD).**

| **Protein** | **Estimate** | **SE** | **p.value** |
| --- | --- | --- | --- |
| GRAP1--GRIPAP1 | -0.698532718 | 0.1737369 | 0.0000580412 |
| FIBG--FGG | 0.574649943 | 0.1635345 | 0.0004415162 |
| CSKI1--CASKIN1 | -0.535646583 | 0.1616413 | 0.0009203780 |
| BSN--BSN | -0.430538522 | 0.1649813 | 0.0090642902 |
| VAMP1--VAMP1 | -0.438276424 | 0.1685093 | 0.0092978458 |
| GSTK1--GSTK1 | -0.417101984 | 0.1607928 | 0.0094857098 |
| OXR1--OXR1 | -0.378492334 | 0.1477391 | 0.0104102369 |
| AHNK--AHNAK | 0.435118202 | 0.1746512 | 0.0127256620 |
| DPP3--DPP3 | -0.417476693 | 0.1712480 | 0.0147749837 |
| KIF5C--KIF5C | -0.379564646 | 0.1586239 | 0.0167176463 |
| PI4KA--PI4KA | -0.419181144 | 0.1770776 | 0.0179223956 |
| SFXN1--SFXN1 | -0.393136704 | 0.1789197 | 0.0280004025 |
| APEX1--APEX1 | 0.411230798 | 0.1908706 | 0.0312009088 |
| DMXL2--DMXL2 | -0.346691503 | 0.1614615 | 0.0317766985 |
| BAG6--BAG6 | 0.373993554 | 0.1829011 | 0.0408760033 |
| SYFB--FARSB | -0.340949880 | 0.1717791 | 0.0471649754 |
| BIG3--ARFGEF3 | -0.276440880 | 0.1426673 | 0.0526645310 |
| SYNPO--SYNPO | -0.315653680 | 0.1664816 | 0.0579563923 |
| HEBP1--HEBP1 | 0.300513978 | 0.1681831 | 0.0739656354 |
| HPRT--HPRT1 | -0.293118532 | 0.1658985 | 0.0772524978 |
| LRP1--LRP1 | 0.304038872 | 0.1728765 | 0.0786275255 |
| CAP2--CAP2 | -0.280350641 | 0.1739887 | 0.1071110817 |
| DHE3--GLUD1 | -0.287873884 | 0.1811871 | 0.1121007283 |
| GNAO--GNAO1 | 0.288843753 | 0.1838025 | 0.1160690211 |
| S12A5--SLC12A5 | 0.273573080 | 0.1842839 | 0.1376710785 |
| SYN3--SYN3 | -0.227969552 | 0.1584390 | 0.1501937664 |
| GRM2--GRM2 | -0.256866010 | 0.1900582 | 0.1765312658 |
| ODO2--DLST | -0.222493023 | 0.1688150 | 0.1875137676 |
| LRC47--LRRC47 | -0.212081725 | 0.1646453 | 0.1977066923 |
| GNAI2--GNAI2 | -0.162150449 | 0.1683075 | 0.3353378187 |
| RAB4A--RAB4A | 0.170805612 | 0.1859644 | 0.3583648258 |
| SYSM--SARS2 | -0.120434472 | 0.1479322 | 0.4155766598 |
| HSP7C--HSPA8 | -0.139017184 | 0.1854018 | 0.4533658064 |
| VGF--VGF | -0.127904783 | 0.1813727 | 0.4806832606 |
| CRKL--CRKL | 0.051351176 | 0.1410139 | 0.7157408471 |
| NRX3A--NRXN3 | -0.034869482 | 0.1453929 | 0.8104625232 |
| GHC1--SLC25A22 | 0.013262041 | 0.1473639 | 0.9282910366 |
| VP26B--VPS26B | -0.003954038 | 0.1695853 | 0.9813982625 |
| MAON--ME3 | 0.001237047 | 0.1448221 | 0.9931846780 |

*These tables report associations between protein abundance and two neuropathological measures of Alzheimer's disease: CERAD scores (reflecting amyloid plaque burden) on top and Braak stage (reflecting neurofibrillary tangle distribution) on bottom. Protein refers to the UniProt identifier and gene symbol. Estimate indicates the effect size from a linear regression model of a dichotomized neuropathology score, adjusted for age, sex, neuron proportion, post-mortem interval. SE is the standard error of the estimate, and p.value is the nominal p-value for the association. Proteins with p < 0.05 were considered nominally significant.*

**Associations between prioritized proteins and neuropathological burden (Braak).**

| Protein | Estimate | SE | p.value |
| --- | --- | --- | --- |
| AHNK--AHNAK | 0.56122791 | 0.2306270 | 0.01495416 |
| GNAO--GNAO1 | 0.57367718 | 0.2398571 | 0.01676844 |
| FIBG--FGG | 0.50635439 | 0.2172025 | 0.01973993 |
| KIF5C--KIF5C | 0.47253748 | 0.2039531 | 0.02050954 |
| GNAI2--GNAI2 | 0.50615295 | 0.2278960 | 0.02635218 |
| LRP1--LRP1 | 0.45996144 | 0.2314766 | 0.04691400 |
| RAB4A--RAB4A | 0.45623090 | 0.2551836 | 0.07379965 |
| CRKL--CRKL | 0.29694336 | 0.1816806 | 0.10216954 |
| GRM2--GRM2 | 0.36094370 | 0.2603946 | 0.16570375 |
| VGF--VGF | 0.29342340 | 0.2315974 | 0.20517152 |
| BAG6--BAG6 | 0.30629560 | 0.2434261 | 0.20829436 |
| HEBP1--HEBP1 | 0.26140928 | 0.2152813 | 0.22464525 |
| PI4KA--PI4KA | -0.24856118 | 0.2291458 | 0.27804163 |
| GRAP1--GRIPAP1 | -0.21316102 | 0.2132710 | 0.31756018 |
| SYSM--SARS2 | 0.18815867 | 0.1974321 | 0.34057504 |
| NRX3A--NRXN3 | 0.17805237 | 0.1870917 | 0.34125681 |
| SYFB--FARSB | -0.19324802 | 0.2116341 | 0.36117778 |
| APEX1--APEX1 | 0.19585859 | 0.2410215 | 0.41643660 |
| CAP2--CAP2 | -0.18848601 | 0.2342474 | 0.42102449 |
| MAON--ME3 | 0.14577112 | 0.1908849 | 0.44506997 |
| SFXN1--SFXN1 | 0.16609876 | 0.2192011 | 0.44860308 |
| HSP7C--HSPA8 | 0.17666801 | 0.2456225 | 0.47197684 |
| DHE3--GLUD1 | -0.16119033 | 0.2336525 | 0.49027457 |
| VAMP1--VAMP1 | -0.13305464 | 0.2156924 | 0.53731895 |
| S12A5--SLC12A5 | 0.14158725 | 0.2323460 | 0.54227190 |
| OXR1--OXR1 | -0.11769150 | 0.1947104 | 0.54554860 |
| VP26B--VPS26B | -0.14049639 | 0.2337415 | 0.54778938 |
| SYNPO--SYNPO | -0.12380398 | 0.2138007 | 0.56254687 |
| ODO2--DLST | -0.13129043 | 0.2308169 | 0.56948668 |
| SYN3--SYN3 | 0.12061399 | 0.2156126 | 0.57588781 |
| BIG3--ARFGEF3 | -0.09515255 | 0.1844750 | 0.60599281 |
| LRC47--LRRC47 | -0.10494445 | 0.2162915 | 0.62753505 |
| DMXL2--DMXL2 | -0.07227735 | 0.2056645 | 0.72526332 |
| GHC1--SLC25A22 | -0.06133509 | 0.1920571 | 0.74945463 |
| DPP3--DPP3 | -0.06552946 | 0.2119331 | 0.75717032 |
| BSN--BSN | 0.06530017 | 0.2230723 | 0.76972786 |
| CSKI1--CASKIN1 | 0.03269136 | 0.1921681 | 0.86491694 |
| GSTK1--GSTK1 | -0.02785325 | 0.1963263 | 0.88718096 |
| HPRT--HPRT1 | -0.02502762 | 0.2194896 | 0.90921682 |

### **Supplementary Table 7**

**Gene Ontology (GO) enrichment analysis of proteins in the APOEe4-relevant network.**

| **GO-term** | **description** | **FDR** |
| --- | --- | --- |
| [GO:0048143](http://amigo.geneontology.org/amigo/term/GO:0048143) | Astrocyte activation | 0.0152 |
| [GO:1900221](http://amigo.geneontology.org/amigo/term/GO:1900221) | Regulation of amyloid-beta clearance | 0.0161 |
| [GO:0032930](http://amigo.geneontology.org/amigo/term/GO:0032930) | Positive regulation of superoxide anion generation | 0.0161 |
| [GO:0016358](http://amigo.geneontology.org/amigo/term/GO:0016358) | Dendrite development | 0.0123 |
| [GO:0007416](http://amigo.geneontology.org/amigo/term/GO:0007416) | Synapse assembly | 0.0123 |
| [GO:0050804](http://amigo.geneontology.org/amigo/term/GO:0050804) | Modulation of chemical synaptic transmission | 0.0062 |
| [GO:0007610](http://amigo.geneontology.org/amigo/term/GO:0007610) | Behavior | 0.0062 |
| [GO:0008088](http://amigo.geneontology.org/amigo/term/GO:0008088) | Axo-dendritic transport | 0.0194 |
| [GO:0050808](http://amigo.geneontology.org/amigo/term/GO:0050808) | Synapse organization | 0.0123 |
| [GO:0060627](http://amigo.geneontology.org/amigo/term/GO:0060627) | Regulation of vesicle-mediated transport | 0.0104 |
| [GO:0099536](http://amigo.geneontology.org/amigo/term/GO:0099536) | Synaptic signaling | 0.0123 |
| [GO:1905908](http://amigo.geneontology.org/amigo/term/GO:1905908) | Positive regulation of amyloid fibril formation | 0.0417 |
| [GO:0006432](http://amigo.geneontology.org/amigo/term/GO:0006432) | phenylalanyl-tRNA aminoacylation | 0.0417 |
| [GO:0002265](http://amigo.geneontology.org/amigo/term/GO:0002265) | Astrocyte activation involved in immune response | 0.0417 |
| [GO:1904589](http://amigo.geneontology.org/amigo/term/GO:1904589) | Regulation of protein import | 0.0427 |
| [GO:1902951](http://amigo.geneontology.org/amigo/term/GO:1902951) | Negative regulation of dendritic spine maintenance | 0.0427 |
| [GO:0007611](http://amigo.geneontology.org/amigo/term/GO:0007611) | Learning or memory | 0.0213 |
| [GO:0070201](http://amigo.geneontology.org/amigo/term/GO:0070201) | Regulation of establishment of protein localization | 0.0161 |
| [GO:0044788](http://amigo.geneontology.org/amigo/term/GO:0044788) | Modulation by host of viral process | 0.0427 |
| [GO:0031175](http://amigo.geneontology.org/amigo/term/GO:0031175) | Neuron projection development | 0.0161 |
| [GO:0006418](http://amigo.geneontology.org/amigo/term/GO:0006418) | tRNA aminoacylation for protein translation | 0.0499 |
| [GO:0032880](http://amigo.geneontology.org/amigo/term/GO:0032880) | Regulation of protein localization | 0.0161 |
| [GO:1901698](http://amigo.geneontology.org/amigo/term/GO:1901698) | Response to nitrogen compound | 0.0161 |
| [GO:0007626](http://amigo.geneontology.org/amigo/term/GO:0007626) | Locomotory behavior | 0.0417 |
| [GO:0071705](http://amigo.geneontology.org/amigo/term/GO:0071705) | Nitrogen compound transport | 0.0123 |
| [GO:0070374](http://amigo.geneontology.org/amigo/term/GO:0070374) | Positive regulation of ERK1 and ERK2 cascade | 0.0427 |
| [GO:0032879](http://amigo.geneontology.org/amigo/term/GO:0032879) | Regulation of localization | 0.0123 |
| [GO:0051049](http://amigo.geneontology.org/amigo/term/GO:0051049) | Regulation of transport | 0.0161 |
| [GO:0051223](http://amigo.geneontology.org/amigo/term/GO:0051223) | Regulation of protein transport | 0.0427 |
| [GO:0007267](http://amigo.geneontology.org/amigo/term/GO:0007267) | Cell-cell signaling | 0.0427 |
| [GO:1901700](http://amigo.geneontology.org/amigo/term/GO:1901700) | Response to oxygen-containing compound | 0.0421 |
| [GO:0042221](http://amigo.geneontology.org/amigo/term/GO:0042221) | Response to chemical | 0.0213 |
| GO:0065008 | Regulation of biological quality | 0.0427 |

*This table presents GO terms significantly enriched among the proteins included in the APOEe4-relevant network. GO-term refers to the identifier of the enriched term; description provides the corresponding biological process name; and FDR indicates the false discovery rate–adjusted p-value for enrichment. GO terms with FDR < 0.05 were considered significantly enriched.*
